# Information Theory Optimization of Signals from Small Angle Scattering Measurements

**DOI:** 10.1101/2024.10.06.616765

**Authors:** Robert P. Rambo, John A. Tainer

## Abstract

Solution-state small angle scattering (SAS) using X-rays (SAXS) or neutrons (SANS), informs on the conformational states and assemblies of biological macromolecules (bioSAS) outside of cryo- and solid-state conditions. BioSAS measurements are resolution-limited and through an inverse Fourier transform, the measured SAS intensities directly relate to the physical space occupied by the particles via the *P*(*r*)-distribution. Yet, this inverse transform of SAS data is canonicaly cast as an ill-posed, ill-conditioned problem requiring an indirect approach. Here, we show that with modern instrumentation and the application of information theory, the inverse transform of SAXS intensity data can be cast as a well-conditioned problem and that the ill-conditioning of the inverse problem is directly related to the Shannon number. By exploiting the oversampling enabled by modern detectors, a direct inverse Fourier transform of the SAXS data is possible provided the recovered information does not exceed the Shannon limit. The limit corresponds to the maximum number of significant singular values that can be recovered in a SAXS experiment suggesting the relationship between the Shannon limit and significant singular values is a fundamental property of band-limited inverse integral transform problems. An intrinsic challenge to the type of problem found in SAS, is identifying the best model while avoiding over-fitting. We propose a hybrid scoring function employing an information theory framework that assesses both the quality of the model-data fit as well as the quality of the recovered *P*(*r*)-distribution. The hybrid score utilizes the Akaike Information Criteria and Durbin-Watson (*D*_*W*_) statistic that considers model complexity, *i*.*e*., degrees-of-freedom and randomness of the model-data residuals. Importantly, the described tests and findings extend the boundaries for bioSAXS by completing the information theory formalism initiated by Peter B. Moore to enable a quantitative measure of resolution in SAS, robustly determine maximum dimension, and more precisely define the best model to appropriately represent the observed scattering data.

**SIGNIFICANCE:** The inverse transform problem in small angle scattering (SAS) of dispersed particles in solution is canonically considered an ill-posed, ill-conditioned problem. We apply information theory to show that the so-called Shannon number, *N*_*s*_, of band-width limited signals defines the boundary at which a problem transitions from well-conditioned to ill-conditioned. In this framework, *N*_*s*_ represents the maximum number of orthogonal elements that are available to the SAS measurement and defines a fundamental relationship that determines the effective resolution of the recovered *P*(*r*)-distribution. We find that the inverse problem can be solved directly using a Shannon-limited approach. The Shannon-limit establishes model complexity and when incorporated into the Akaike Information Criteria aids in selecting the most likely *P*(*r*)-distribution.

## INTRODUCTION

Small angle scattering of X-rays (SAXS) and neutrons (SANS) by particles in solution is an established technique for investigating the conformational and assembly states of biological macromolecules outside of cryo- and solid-state conditions (1). A bioSAS measurement results from the scattering of billions of randomly oriented molecules providing statistical power to single particle, microscopic observations. Under sufficiently dilute conditions, the bioSAS experiment represents a resolution-limited measurement of the particles’ thermodynamic solution-state. It is a measurement that directly informs on the physical space occupied by particles in solution and can be envisaged as a weighted average of the thermodynamic ensemble (2).

A SAXS dataset consists of a set of measured intensities, *I*(*q*), defined within a scattering vector (*q*) range [*q*_*min*_, *q*_*max*_]. Here, the scattering vector, *q*, is 4*π* · *sin*(*θ*) · *λ*^−1^ in units of reciprocal Å with *θ* as the half-scattering angle and *λ* as the X-ray wavelength in Å. The scattering vector range is a consequence of the instrument’s geometry that includes sample-to-detector distance, active detector area, beamstop size, and X-ray source properties (*e*.*g*., divergence, wavelength). The useable scattering vector range is often a subset of the instruments’ observable range where the particle scattering intensity must be sufficiently greater than the parasitic scattering from the beamstop, slits and sample cell windows to optimally define *q*_*min*_. At larger *q*-values, the quality of the buffer matching (background subtraction) and particle concentration will be major factors that determine *I*(*q*_*max*_). The useable [*q*_*min*_, *q*_*max*_] range defines the set of intensities that faithfully describes the scattering of the particles under investigation with minimal contributions from parasitic scatter and poor buffer matching.

Importantly, the useable SAXS dataset will consist of *J* points within [*q*_*min*_, *q*_*max*_]. This number has steadily increased with improvements in detector technologies from a few hundred points using a CCD detector to over 2400 points with the latest generation photon counting detectors. The increased number of points within a fixed [*q*_*min*_, *q*_*max*_] implies a greater over-sampling of the underlying SAXS signal. If we consider a fixed [*q*_*min*_, *q*_*max*_] bioSAXS measurement that measures a 21 kDa particle at 1 mg/ml using a detector with 200 points versus 2000 points, the information content obtained from each measurement will be the same assuming the same point focused camera geometry. However, fitting the same incorrect model to both datasets will demonstrate that the dataset with the greater number of points will artificially drive down a scoring metric such as chi-squared, *χ*^2^, thus contributing to a fundamental false-positive problem in SAXS (3). This observation establishes a paradox in bioSAXS where improvements in detector technologies that decrease pixel sizes will improve model-data agreements regardless of the model when using scoring metrics that treat each individual *I*(*q*) as an independent observation.

A critical path to resolving this conflict was presented by Peter B. Moore (4) in 1980. He presented a formalism of SAXS based on information theory (5) which demonstrated that the number of independent data points, *N*_*s*_, that could be measured up to a given *q*_*max*_ can be estimated as *N*_*s*_ = *q*_*max*_ · *d*_*max*_ · *π*^−1^. For a particle with a *d*_*max*_ of 240 Å measured to *q*_*max*_ of 0.4 Å^-1^, implies ∼31 data points are required to fully capture the information within the scattering vector range measured at 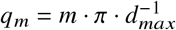 Furthermore, Moore showed that each measured intensity, *I*(*q*_*j*_), is a linear combination of a fundamental set of intensities measured at *I*(*q*_*m*_) (see SI). This observation implies modern SAXS datasets are highly redundant and correlated (3).

By using the information theory framework, we previously showed (6) that application of the Shannon-Hartley theorem (noisy-channel coding theory) (7) guarantees error-free recovery of the SAXS signal as long as the sampling frequency, *i*.*e*., distance between successive *q*-values, is within the Shannon-Hartley limit (see SI). However, this finding imposes a fundamental question: what exactly is the signal that is being sampled by SAXS? Based on the classical description of SAXS in equation 1 (Eq. 1), an intensity, *I*(*q*), is a sine-integral transform (see SI) of the paired-length, *P*(*r*), -distribution function suggesting the measured SAXS signal is a resolution-limited sampling of the *P*(*r*)-distribution. The integral transform is defined within the maximum dimension (*d*_*max*_) of the particle, however, we take *d*_*max*_ to be defined within the structural space of the measured thermodynamic ensemble.

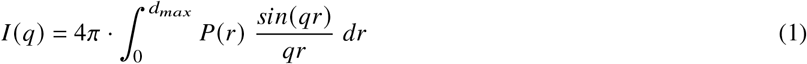

Building upon the above considerations, we show here that information theory provides a complete theoretical framework to enable a direct inverse Fourier transform of SAXS data to accurately recover the *P*(*r*)-distribution. Until now, the inverse transform has been canonically cast as an ill-posed, ill-conditioned (8, 9) problem; however, we show that with modern instrumentation and the application of information theory, the problem is in fact not necessarily ill-posed or ill-conditioned. This formalism allows the direct inverse Fourier transform and provides an intuitive description of resolution in SAXS. Moreover, we provide a general method for determination of the useable scattering range [*q*_*min*_, *q*_*max*_] from a SAXS measurement that addresses the quality of the background subtraction. This report provides the primary literature reference for these insights and algorithms, which are implemented in the program ScÅtter (https://github.com/rambor/scatterIV) for free access by the scientific community. We expect these algorithms and foundational concepts will provide new insights for the understanding of experimental results from ongoing and advanced X-ray and neutron scattering analyses.

## MATERIALS AND METHODS

BMV RNA and xylanase samples were manually purified using a KW 402.5 (Shodex) column in buffer containing 20 mM MOPS at pH 6.5, 50 mM KCl, and either 7.6 mM MgCl_2_. Peak fractions were taken for SAXS at SIBYLS beamline 12.3.1 (Advanced Light Source, Berkeley, CA). BSA samples were prepared at beamline B21 (Diamond Light Source Ltd, Didcot UK). 10 mg of BSA was diluted to 5 mg per ml in PBS buffer. For SEC-SAXS, 45 uL sample was injected on to a KW-403 (Shodex) column at 160 uL per min. SAXS data was collected at 2 second intervals for 32 minutes. In all cases, an X-ray wavelength of 1 Å was used during measurements.

SAXS datasets were reduced with DAWN (B21) and in-house software (SIBYLS). All datasets were processed using the program ScÅtter (https://github.com/rambor/scatterIV).

For the model selection search, each model is scored according to Eq. 10 and sorted. The lowest score of the set is then selected as the best model and assigned an initial probability of 1. For each additional model in the set, a probability is assigned based on how different the score, s, is from the best using Eq. 2

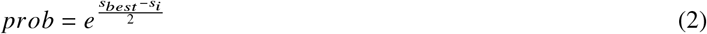

The algorithms for the methods described above (see SI) are coded in the program ScÅtter (JAVA 1.8) available at https://bl1231.als.lbl.gov/scatter/ and the source code can be viewed at the github repository (rambor/scatter3) under src/Version3/InverseTransform.

Since we are interested in the most likely *d*_*max*_, which will be tested through multiple *α* values, we sum the results from Eq. 2 at constant *d*_*max*_ for each *α* to give an amplitude *G*. The scores are subsequently normalized across the entire set and a Gaussian kernel density estimation, KDE, is performed using a window of 2.5. The KDE represents the final likelihood score in Figure 6. For each *d*_*max*_ in the search space, we calculate the following KDE

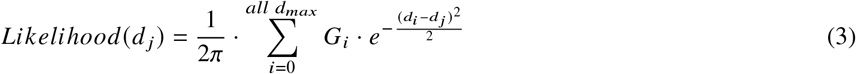

## THEORY AND RESULTS

The Debye equation (Eq. 1) relates the X-ray scattering intensity, *I* (*q*), to the *P*(*r*)-distribution (10). The relationship in Eq. 1 suggests the *P*(*r*)-distribution can be recovered from the SAXS intensities through an inverse integral transform. Yet, in practice the results of the inverse transform are often undesirable due to variances in *I* (*q*), the sampling frequency of the observed *q*-range (Δ*q*) and the choice of *r*-values. It has been proposed that the inverse transform could be performed using an orthogonal set of basis functions such as cubic splines(8), sine function (4), Chebyshev or Zernicke (11, 12) polynomials such that a least squares minimization could be used to perform the inverse transform through an indirect method. Alternatively, the relationship in Eq. 1 can be described as a matrix operation (Eq. 4)

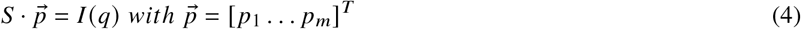

where the matrix, ***S***, contains the elements of the sine transform and the vector 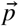 contains the set of unknown values of the *P*(*r*)-distribution (see SI). For a set of *J* intensities and *M* equally spaced points along the domain of the *P*(*r*)-distribution, *S* has the dimensions of *J* rows and *M* columns.

If the errors in the measurements or the number of elements in 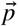 is too large, the problem is ill-posed and the matrix *S* is ill-conditioned (13) thereby preventing a sensible linear least squares solution. In these cases, a solution can be found by imposing expectations (e.g., positivity or smoothness) upon 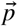 through a technique known as regularization (9, 14).

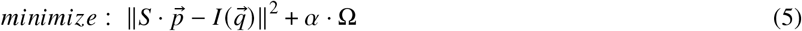

Here, an objective function is defined that seeks to minimise the difference between the calculated, *S* 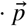 and observed, 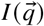, intensities. The objective is augmented with the regularization function, Ω, which embodies the expectation and the influence of Ω is controlled by *α* (*α* ≥ 0). Notably, as *α* tends to infinity, the elements of 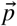 will tend to zero. Therefore, a proper *α* value must be determined empirically (8, 9, 14).

The popular programs GNOM (9) and BayesApp (14) implement a smoothness expectation for recovering the *P*(*r*)-distribution from SAXS intensities. Smoothness can be estimated from the second derivative of the *P*(*r*)-distribution, and in the case of GNOM, Ω is simply the squared sum of the second derivative evaluated at the points defined in 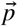. The squared sum of a vector is a length metric that is strictly positive and commonly referred to as the L2-norm. Alternatively, the magnitude of the expectation vector could be evaluated as the sum of the absolute values, a metric known as the L1-norm. The L1-norm is statistically robust and seeks to find a set of parameters that is least. In this regard, regularization using the L1-norm can be considered the least biased estimate of 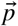 (15). Fundamentally, how many *r*-values should be used to construct 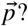 If one uses an orthogonal basis set to represent the *P*(*r*)-distribution, then how many terms should be included in the expansion? An arbitrarily large number of points or terms leads to an ill-conditioned problem. We can determine the condition number of the problem by calculating the singular value decomposition (SVD) of the matrix *S*. Here, the Debye equation is rearranged as the Fourier sine transform as in (4) where the integral transform is first re-arranged to remove the constant term from the right-hand side (rhs).

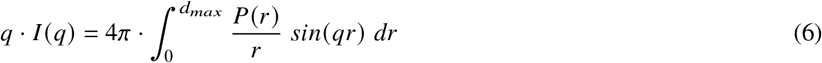

If we let *U*(*q*) = *q* · *I* (*q*) and *Q*(*r*) = *P*(*r*) · *r* ^−1^ then the Debye equation is put into standard form illustrating the Fourier sine integral transform relationship between intensity space, *U*(*q*), and real-space, *Q*(*r*).

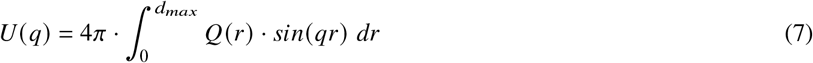

The *M* unknowns are embodied by *Q*(*r*). Each term of *S* is therefore, *sin*(*q* _*j*_ · *r*_*m*_), where *j* indexes over the observed [*q*_*min*_, *q*_*max*_] and *m* indexes over an arbitrary number of equally spaced *r*-values within [0, *d*_*max*_]. From Eq. 4, the unknown terms in 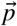 could be recovered if *S* was invertible. It can be expected that *S* will not be a square matrix, *i*.*e*., *M* = *J*, and likewise not invertible. For non-square matrices, a pseudo-inverse, *S*^+^, can be calculated based on an SVD factorisation of the *S*-matrix.

SVD of *S* will produce *K* singular values, *σ*_*k*_, where *K* = *min*(*J, M*). Singular values are arranged from largest to smallest and *S*^+^ can be calculated by setting all insignificant singular values to zero. This effectively reduces the matrix to a set of linearly independent columns. A judgement must be made as to when a *σ*_*k*_ is insignificant. We can quantify this by noting that the condition number of a matrix is the ratio of the first singular value, *σ*_1_, to the last. A well conditioned problem will have a low condition number, *κ*, ideally equal to unity. By plotting the ratio of *σ*_1_ to all subsequent singular values in the set, we can demonstrate how the conditioning of the matrix evolves as the number of unknowns increase through the size of 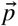.

Using several SAXS datasets of particles with varying *q*_*max*_ and *d*_*max*_ values (Figure 1A and B), the *S*-matrix was calculated for an arbitrary number of *r*-values and an SVD was performed to determine the singular values for each *q*_*max*_, *d*_*max*_ pair (referred to as [*q, d*]_*max*_). Plotting the ratio of the singular values, as described above, shows a general trend where there exists a region of small condition numbers that abruptly increase (Figure 1C). The magnificent increase to infinity for each *S*-matrix illustrates the ill-conditioned behaviour for each combination of *q*_*max*_ and *d*_*max*_. The abrupt transition demarcates the boundary where all subsequent singular values can be considered insignificant as the division by an exceedingly small number drives the ratio to infinity. Interestingly, this boundary is coincident with *N*_*s*_, the Shannon number derived by Moore and suggests *N*_*s*_ can be used to determine the number of significant singular values for a given [*q, d*]_*max*_.

**Figure 1:**
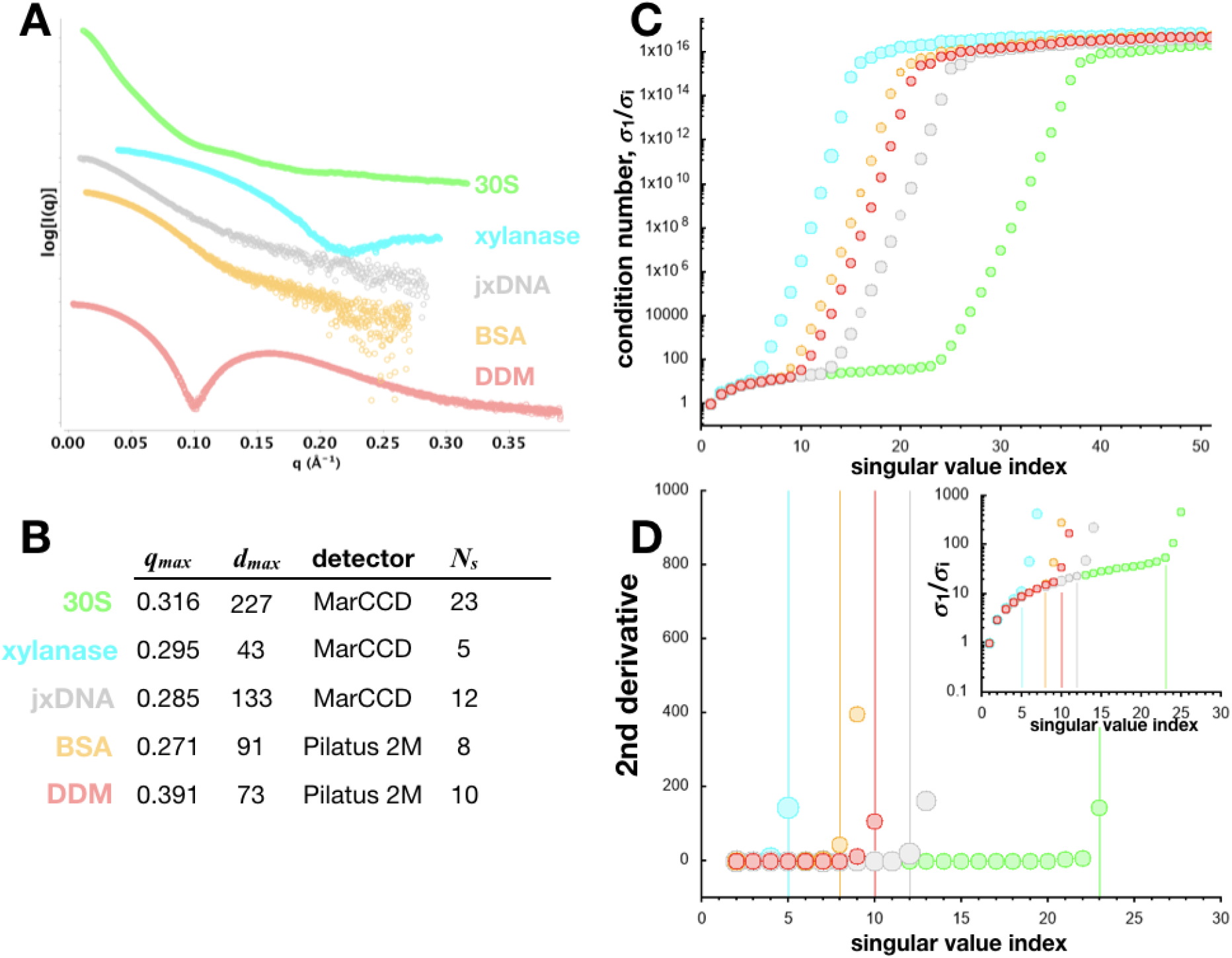
Exemplary SAXS datasets of particles with varying size, shape and composition. A) Experimental datasets for the *Sulfolobus solfataricus* 30S ribosomal subunit (green), *B. subtilis* xylanase (cyan), 25 base-paired double-stranded DNA with 11 nucleotide overhang (gray), Bovine serum albumin, BSA (orange) and n-Dodecyl-beta-D-maltopyranoside, DDM, micelle (red). 30S, xylanase and jxDNA samples are size-exclusion purified samples collected as a batch-mode SAXS experiment (16, 17). BSA and DDM were collected as SEC-SAXS experiments in PBS buffer. B) SAXS datasets were collected on either a CCD detector(18) or a silicon photon counting detector (Pilatus 2M) with a delta-*q* spacing of 0.000608 and 0.0002749 Å^-1^, respectively. Each dataset contains N data points with different [*q, d*]_*max*_ and different Shannon numbers, *N*_*s*_. C) For each [*q, d*]_*max*_, singular value decompositions were performed on *S*-matrices with dimensions of *N* rows and 11×*N*_*s*_ columns. For each *S*-matrix, condition number plots were constructed by plotting the ratio of the i^th^ singular value to the first singular value. Insignificant singular values will drive successive condition numbers several orders of magnitude (note, y-axis is log_10_). The upper plateau is an artefact of floating point machine precision. D) Second derivative plot of the starting data in C shows condition number rate of change for each dataset in A. All datasets illustrate a second derivative that has an initial constant region followed by an abrupt increase. Shannon number for each dataset in B is indicated by the corresponding vertical line. Inset is the condition number plot of the subset of the data in D. DDM dataset was kindly provided by Mark Tully, ESRF (Grenoble, FR).

If the *S*-matrix is constructed using only *N*_*s*_ columns, then the number of unknowns will be restricted to the number of significant eignevalues providing for a well-conditioned problem. Since the integral transform in Eq. 7 are only defined between 0 and *d*_*max*_, there will be a set of *r*-values separated by a Δ*r* of *d*_*max*_/*N*_*s*_ for each of the *sin*(*q* _*j*_ · *r*_*m*_) terms in the *S*-matrix. Here, the *P*(*r*)-distribution is conceptualised as a histogram that is divided into *N*_*s*_ bins where each bin has equal bin-widths of Δ*r*. The *p*-vector is therefore the height of each bin. Fig 2 shows results for several SAXS datasets that are solved for the corresponding 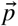 using the pseudo-inverse, *S*^+^.

**Figure 2:**
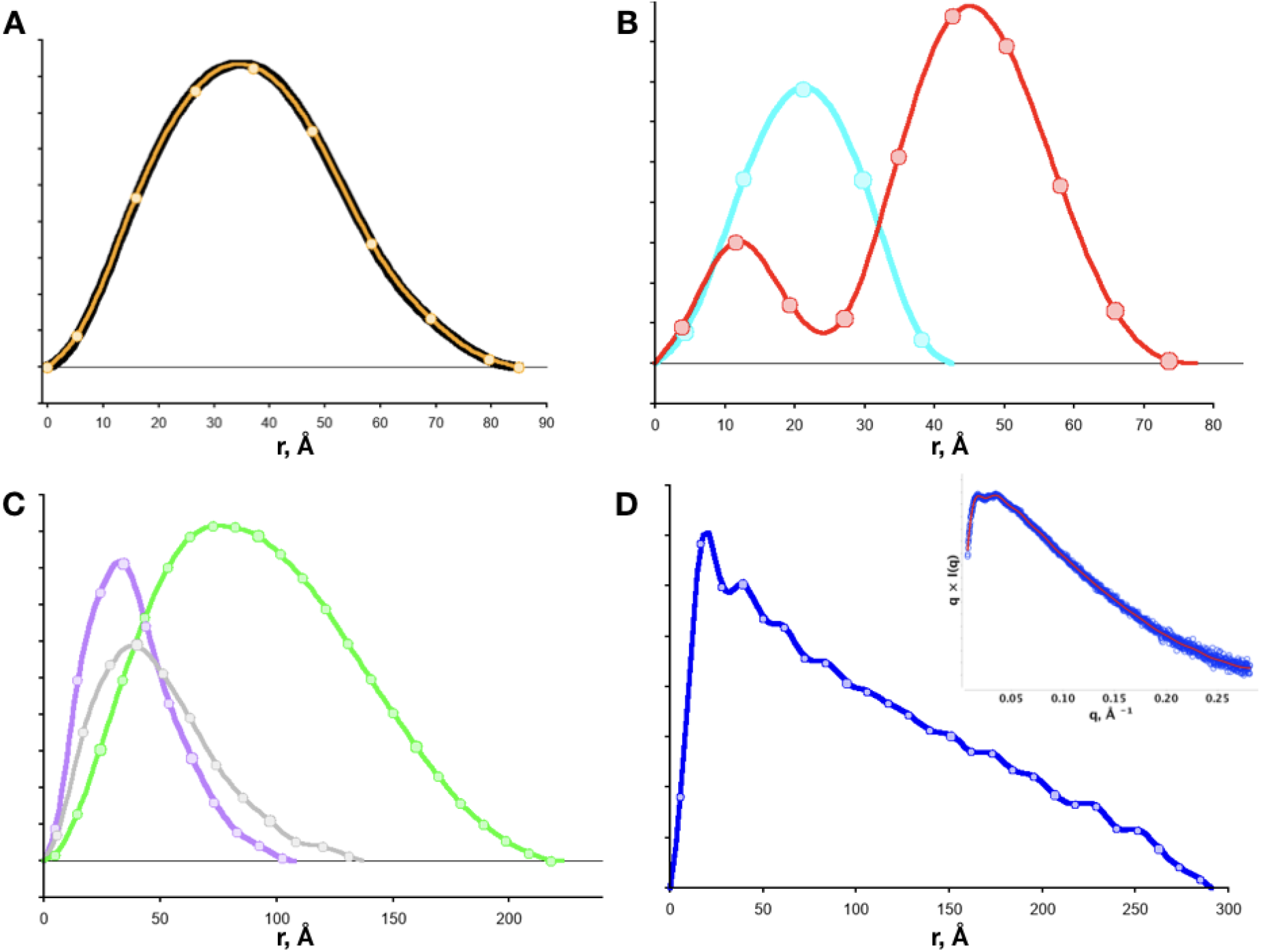
Recovery of *P*(*r*)-distributions using the Shannon-limited, pseudo-inverse (SPI) method for data in Figure 1. A) Transformation of the BSA data using GNOM (black) (9) and SPI (orange). Orange curve is the spline interpolated curve based on the SPI determined points (circles). B) Transformation of xylanase (cyan) and DDM (red) C) SAXS data of nucleic acid proteins from Figure 1. D) SPI-method applied to SAXS data (inset, plotted as *q* × *I* (*q*) vs *q*) of SEC-SAXS data of KASH5 coiled-coil protein. KASH5 data was kindly provided by Owen Davies, New Castle University. Y-axis is in relative units, distributions in B and C are scaled arbitrarily for presentation.

Monomeric BSA with a [0.277174, 85]_max_ corresponds to an *N*_*s*_ of eight which implies an *S*-matrix of eight columns and a 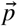 with eight unknowns. The direct recovery of the *P*(*r*)-distribution shows a 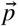 consistent with a distribution (Figure 2A). In fact, overlaying the components of 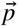 on to an L2-regularized indirect inverse transform that used 246 points shows an ideal coincidence (Figure 2A). Furthermore, a spline interpolation based on *p*-shows a nearly perfect overlay with the L2-regularized inverse transform (Figure 2A) suggesting the Shannon-limited, pseudo-inverse (SPI) method recovers the same information as the L2-regularization method.

The SPI method was applied to additional SAXS datasets (Figure 2) of particles with varying aspect ratios and [*q, d*]_*max*_ pairs. The globular particles xylanase and BSA produced the expected bell-shaped distribution with xylanase requiring only 5 *N*_*s*_ points to represent 496 reciprocal-space data points. We also applied our method to mixed phase particles such as a DDM micelle and the 30S ribosomal subunit. For the DDM micelle, the recovered 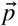 with 10 *N*_*s*_ displays the classic bimodal distribution of detergent micelles whereas the 30S subunit at a *d*_*max*_ of 227 Å requires 23 *N*_*s*_ points to describe the large, asymmetric particle. Finally, we examined three highly asymmetric particles as the SPI method should be insensitive to particle shapes. The P4-P6 RNA domain, a 25 base-paired (bp) double-stranded (ds)DNA with 11 nucleotide overhang and a coiled-coil protein domain with a *d*_*max*_ larger than the 30S subunit were examined. In all cases, the recovered 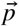 produced a smooth distribution, however, the coiled-coil SAXS dataset shows subtle oscillatory features that could be real or an aliasing artefact. Oscillatory features can result from special conditions between the interpolating polynomial and the underlying equispaced points (Runge’s phenomenon (19)). It must be emphasised that the equispaced points from the SPI method represent the actual resolution-limited, real-space data from the SAXS measurement whereas the oscillation is a visual artefact. As the number of real-space points increases or decreases, oscillatory features from special conditions will vary, so a completely smooth, oscillation-free *P*(*r*)-distribution should not be a necessary condition for an acceptable *P*(*r*)-distribution. This can be best appreciated by examining *q*_*max*_-dependent changes in the atomistic *P*(*r*)-distribution calculated from a 137-bp dsDNA model (Fig. S1A). At a *q*_*max*_ of 0.302 Å^-1^, there are 14 *N*_*s*_ points that provide a relatively coarse description of a skewed distribution and as *q*_*max*_ increases, the number of bins that define the *P*(*r*)-distribution will increase producing a finer description of the distribution. More importantly, at a *q*_*max*_ of 0.302 Å-1, the bin-width matches the pitch (10.4 Å) of an B-form dsDNA helix and we see at this resolution, the *P*(*r*)-distribution is exceptionally smooth. However, at half-integer increments of 0.302, the distribution develops oscillatory features that appears to be most severe at even number increments of 0.302 and least at odd number increments. The oscillation is due to an undulation between successive bins and since the bin heights are calculated directly from the structure, the oscillation is entirely a consequence of the bin-width and does not represent an unstable solution of the inverse transform.

Regularization is a widely used tool for performing the inverse transform of SAXS data (8, 9, 14). It imposes an expectation on 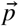 and in the case of smoothness, the regularisation attempts to minimise the differences between neighbouring values in the *P*(*r*)-distribution. Yet as discussed above (Fig. S1A), such an expectation may unduly attenuate real features in a distribution as smoothness becomes an over-valued expectation, *i*.*e*., large *α*. Nevertheless, regularisation has been essential in transforming smeared, noisy and under-sampled SAXS data. Todays SAXS measurements will mainly suffer from noise, particularly as measurements are made at low particle concentrations and regularisation may be critical to a successful inverse transform. The determination of *S*^+^ via SVD does not consider noise in the dataset. Figure 3 shows application of the SPI method on SAXS data with a large variance between successive data points (Figure 3B).

**Figure 3:**
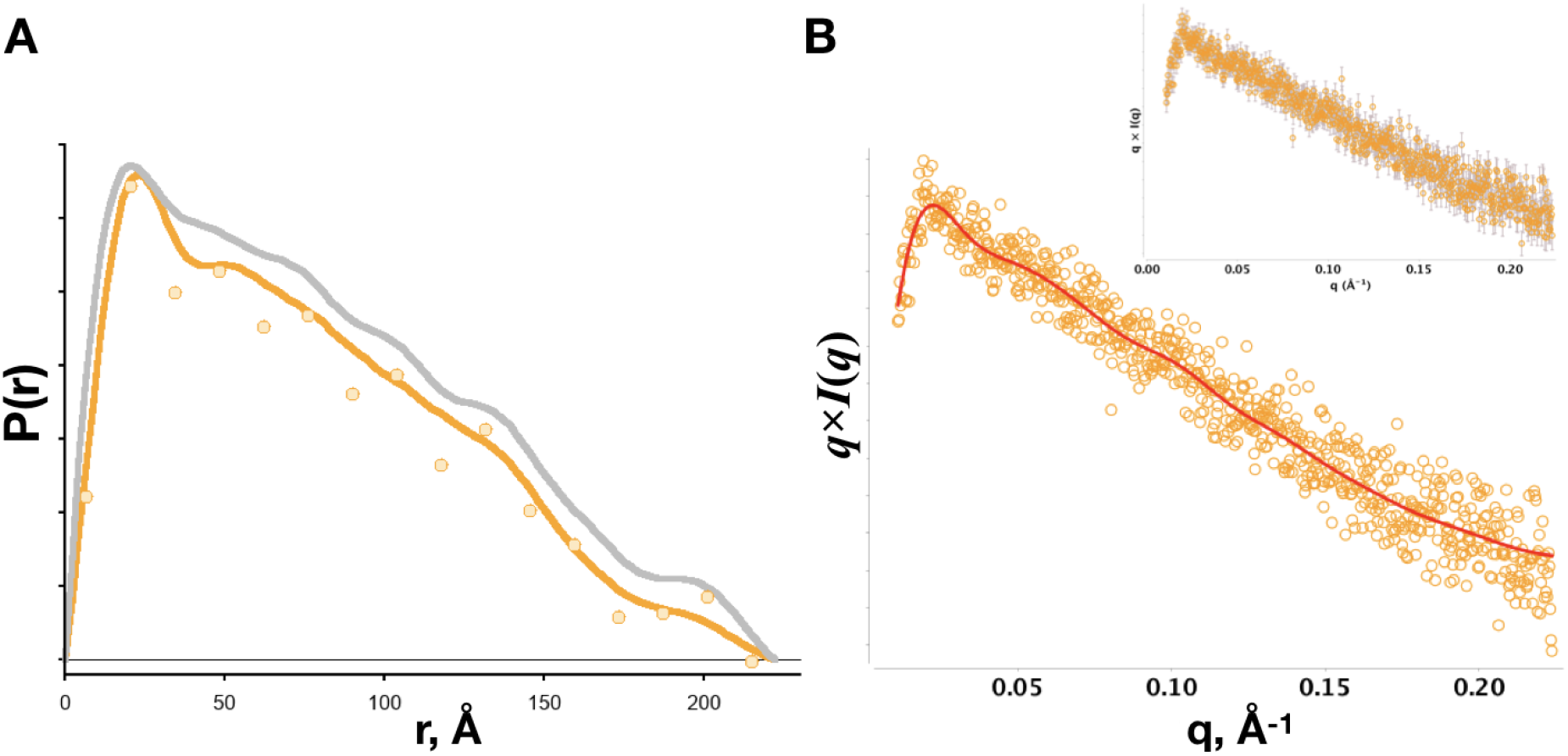
Real-space transformation of a challenging, high noise SAXS dataset from a high aspect ratio particle (the coiled-coil protein SYP1 pH 8, (20)). A) *P*(*r*)-distribution recovered using inverse transform methods GNOM (gray line), SPI (orange circle) and Moore (orange line) with smoothness regularization as in GNOM. Using an excessive number of points (GNOM default 256 points) to describe the distribution has the effect of filling in features as smoothness minimizes differences between neighboring points (see valley near 40 Å). Y-axis plots the distribution in relative units. The data was kindly provided by Owen Davies, Institute Cell and Molecular Biology Newcastle University, UK. B) SAXS dataset transformed as *q* × *I* (*q*) with the fit from the Moore method in A (red line). Inset shows same data with error bars.

The SPI method fails to produce a solution that is smooth, the oscillations persist through several choices of [*q, d*]_*max*_. Here, regularisation will be critical to finding an appropriate solution. However, there will be variations in the solution depending on how the regularised inverse transform is performed (Figure 3A and SI Fig 2), *i*.*e*., the size of *p*- and the nature of the regularisation. The goal is to recover the real-space information that is least biased.

The condition number analysis (Figure 1) suggests *N*_*s*_ determines the maximum number of linearly independent columns of the *S*-matrix. Columns that are linearly independent are orthogonal which implies the *P*(*r*)-distribution can be represented as a set of orthogonal functions (21). In fact, the orthogonal series expansion is the basis for the methods proposed by Glatter (8) and Moore (4). In Glatter’s indirect Fourier transform (IFT) method, the distribution is approximated as a set of cubic B splines (orthogonal polynomials) whereas in Moore’s method, the distribution is approximated as a Fourier sine series (orthogonal trigonometry functions). In addition, the IFT implements a smoothness-based regularisation. However, since the *P*(*r*)-distribution is strictly positive and only non-zero within a defined domain [0, *d*_*max*_], we can alternatively represent the *P*(*r*)-distribution using the orthogonal set of Legendre polynomials, *L* (*r*). Legendre polynomial density approximations are inherently smooth and provide a well established technique that makes no assumptions on the shape of the distribution (22).

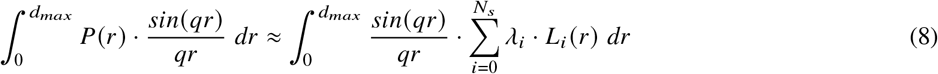

In this approach, each element in the *S*-matrix will be a product-sum of the sine term and a Legendre polynomial whereas *p*-contains the set of unknowns, *λ*_*i*_, that can be determined through least-squares (see SI). To test the robustness of the *N*_*s*_ limited polynomial expansion fitting of SAXS data, we examined the coiled-coil data in Figure 1D and a dataset of unpurified BSA in PBS buffer (Figure 4). The coiled-coil protein has a high aspect ratio with a cross-sectional diameter of 12 Å and a length of nearly 300 Å whereas the BSA dataset will be of a mixture of monomers and dimers with some higher order oligomeric species suggesting the solution-state ensemble will consist of particles of varying *d*_*max*_ values. In comparison to the SPI method, the Legendre polynomial expansion method resulted in smoother distributions that are nearly identical at the extremes but differ within the medium range distances for both datasets (Figure 4). Furthermore, the *N*_*s*_ limited polynomial expansion was tested using the Moore method (Figure S2). The recovered Fourier sine series distribution showed substantial agreement with the Legendre-based method, however there are subtle differences that are likely related to the type of basis function used in the expansion.

**Figure 4:**
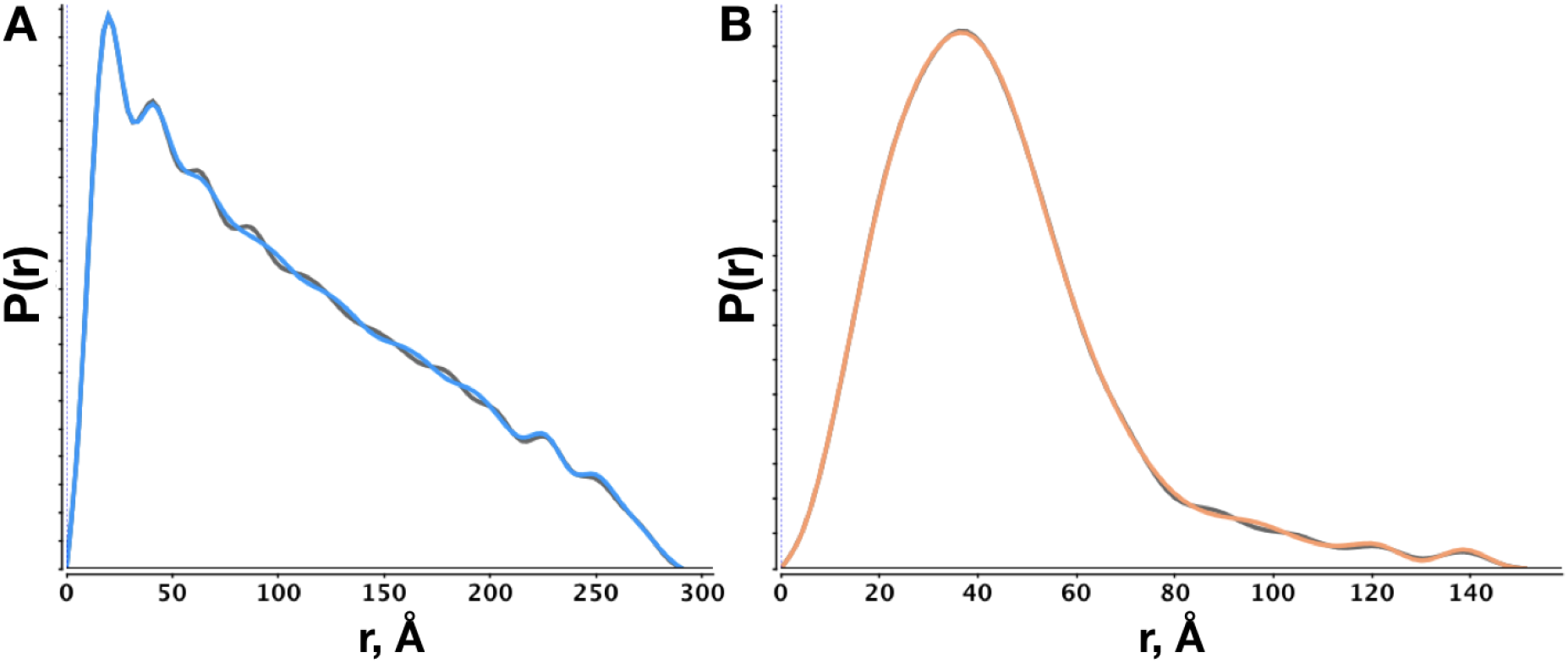
Indirect Fourier transforms (IFTs) using an orthogonal series expansion without regularization (*α* = 0) A) IFT of coiled-coil data in Figure 1D using Legendre (blue) and SPI (dark gray) methods. B) IFT of unpurified BSA using Legendre (orange) and SPI (dark grey) methods. *D*_*max*_ was fixed to the same value for each method in the respective datasets.

Increasing the number of expansion terms beyond *N*_*s*_ introduces instabilities or artefacts in the recovered distribution as the inverse transform problem becomes ill-conditioned. For both the Moore and Legendre-based methods, instabilities will contribute to oscillations and spurious features in the recovered distribution (Figure S3 and S4). For a given [*q, d*]_*max*_, profiling the condition number (Figure 1) shows how sensitive the *S*-matrix will be to the number of unknowns, *M*. By restricting the matrix inversion or orthogonal series expansion to *N*_*s*_, we maximize the information recovered while minimising artefacts introduced from regularisation or ill-conditioning.

The Shannon number is a quantitative measure that informs on the number of independent columns of the *S*-matrix or terms used in an orthogonal series expansion. It represents the maximum number of non-degenerate elements of information contained within the solution-state SAXS measurement defined by [*q, d*]_*max*_. Practically, both *q*_*max*_ and *d*_*max*_ are unknowns in a SAXS measurement as the former is not solely determined by the physical limitations of the detector. The largest scattering vector that contains useable intensity information, *I* (*q*_*max*_), cannot be determined simply by a signal-to-noise ratio, *I* (*q*_*i*_)/*σ*_*i*_, as the Shannon-Hartley theorem stipulates that the ability to recover a signal relies on both the magnitude of *I* (*q*_*i*_)/*σ*_*i*_ and the sampling frequency, Δ*q*. Likewise, assessing the quality of the inverse transform using a chi-squared metric alone has proven not to be reliable for choosing the appropriate *d*_*max*_ due to over-fitting (4, 6, 9, 14). The assessment must consider both the quality of the model-data agreement and the quality of the putative *P*(*r*)-distribution.

In practice, determining both *q*_*max*_ and *d*_*max*_ is an iterative process where the recovered *P*(*r*)-distribution is visually assessed for smoothness, non-negativity and a shallow finish. If *q*_*max*_ is too large, the dataset may include intensities that carry no information. Here, the buffer subtracted intensities at high-*q* will be randomly distributed around some average value, *b*. Ideally, the average should be zero, but for imperfect buffer subtracted samples, *b* will be a non-zero constant that the SAXS intensities at high-*q* will be distributed around. The effect of *b* can be readily illustrated using SEC-SAXS data of BSA collected under very dilute conditions (0.26 mg/ml). The sample is ideally buffer matched however, the intensities become distributed near zero (Figure 5A) beyond moderate scattering vectors (*q* > 0.2). Performing the inverse transform of the SAXS data without a constant background term resulted in a bulging of the distribution near *d*_*max*_ (Figure 5B) using the Legendre, Moore and SPI methods. This bulging persisted with a SAXS dataset of the same sample that was averaged over 17 independent SEC-SAXS runs suggesting the low signal-to-noise ratio is not responsible for the bulge. In addition, attenuating the bulge through a large *α* introduced artefacts in the distribution or led to an overestimated *d*_*max*_. For both SAXS datasets, performing the inverse transform with a constant background term in the *S*-matrix provided for an ideal *P*(*r*)-distribution (Figure 5C and S5). Notably, regularisation was critical to the successful inverse transform suggesting that for weakly measured signals near or within the background, the regularisation with a constant *b* term can optimally recover a *P*(*r*)-distribution. The contribution of *b* becomes significant under very dilute conditions as the subtracted intensities are within the background. The q-value at which the intensities become evenly and randomly distributed about *b* suggests a *q*_*max*_ for the SAXS dataset.

**Figure 5:**
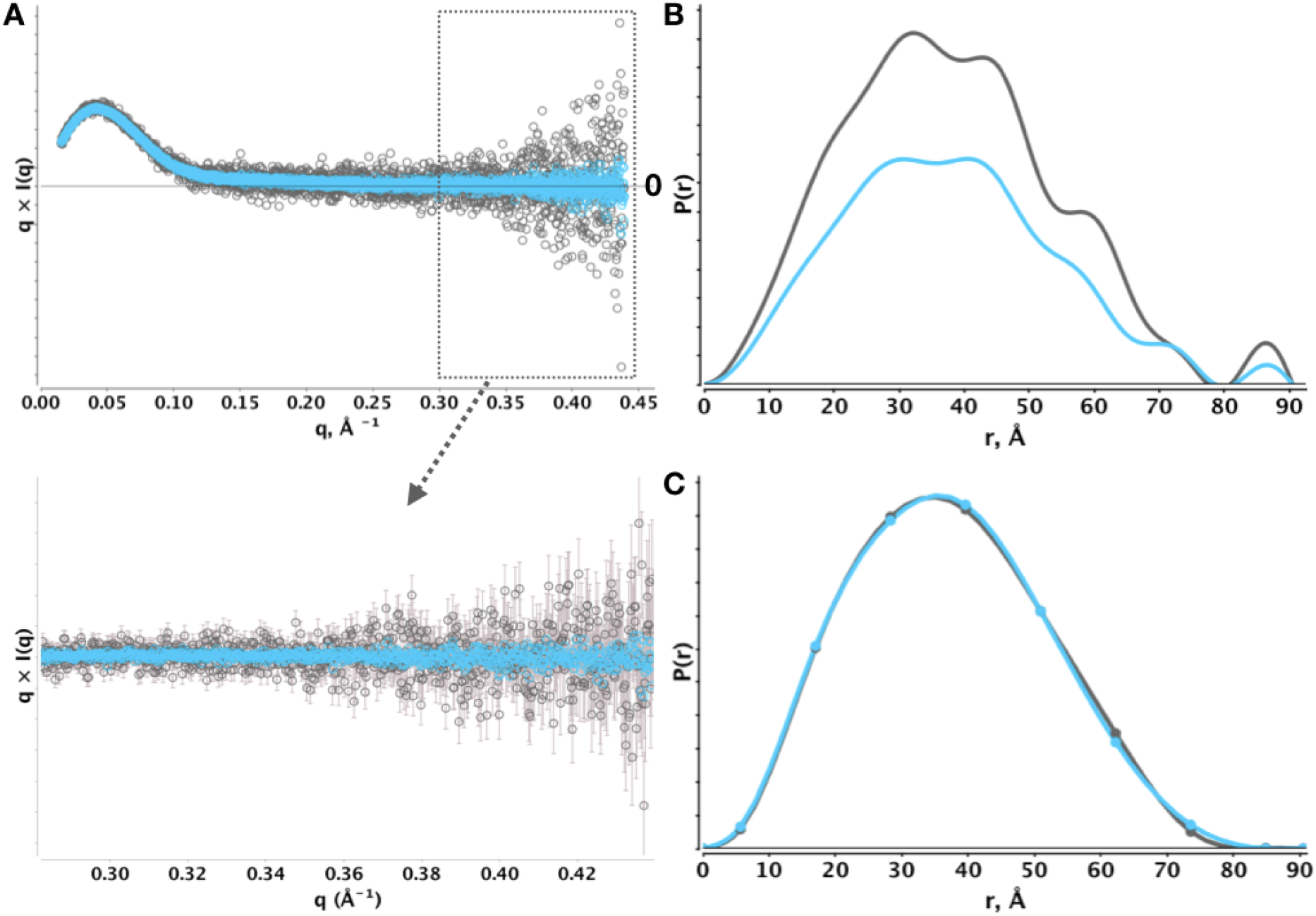
Monomeric BSA from SEC-SAXS of dilute BSA in PBS buffer. A) Average of ∼20 frames across a single elution peak (dark gray circles) corresponding to ∼0.26 mg per ml versus an average of 18 independent, identical SEC-SAXS runs of monomeric BSA at the same ∼0.26 mg per ml concentration. Lower left panel (arrow) is subset of data plotted above with errors. B) L1-norm regularized SPI method for both datasets in A. C) Regularized SPI method with constant background term as in B. For both B and C IFT was achieved with *α* = 0.007, *d*_*max*_ 90.5 Å.

## Model selection

A given [*q, d*]_*max*_ and *α* collectively define the parameters of a particular inverse transform model (e.g. Moore, Glatter, SPI). Therefore, finding the “best” model is a search in a multi-dimensional parameter space where each model must be evaluated not only for the quality of the recovered *P*(*r*)-distribution, Ψ, but also for the quality of the fit to the experimental intensities. Such a search represents a model selection within a constrained search space and since the complexity of the model will scale with *N*_*s*_, the selection of the “best” model must weigh the risk of over-fitting. This can be readily achieved by incorporating the Akaike information criterion (23) into an integrated score (24). AIC has been successfully applied to ensemble modeling of SAXS data using atomistic models (24). Here, we use a modification of the criterion (*AIC*_*c*_) that includes corrections for small samples sizes that are imposed by *χ*^2^-*free* (6).

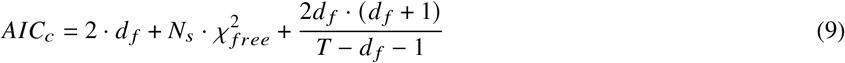

where *d* _*f*_ is the degrees-of-freedom and *T* is the sample size used to calculate *χ*^2^-*free*. The *d* _*f*_ is taken as *N*_*s*_ plus three that includes assumptions on *α, d*_*max*_ and that the residuals between the model-data agreement are Gaussian. To assess the model-data agreement, we will use an alternate to *χ*^2^ by noting that an ideal agreement is characterised by a random distribution of the residuals. We quantitate this randomness via the Durbin-Watson statistic (D_W_) which examines the auto-correlation of the residuals with a lag of 1 (25). D_W_ is bounded between 0 and 4 and will be 2 for perfectly random residuals. Thus, we can construct a metric as the absolute value of (2 -D_W_). For Ψ, we require a function that quantitates the finish near *d*_*max*_, the overall smoothness of the *P*(*r*)-distribution and penalises for negative values [see SI]. Our overall score is given by Eq. 10

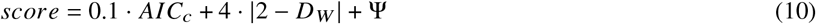

with the scaling factors empirical determined. At fixed *q*_*max*_, our scoring function can be applied to a set of models that vary by *d*_*max*_ and *α* where the minimum score identifies the best model of the set. However, we can collectively combine the scores into a likelihood function that identifies the region of the most likely *d*_*max*_ [see SI].

Using SEC-SAXS data of a coil-coiled and BSA (Figure 5A and B), the scoring function shows a clear minimum at the expected *d*_*max*_. The scoring function is relatively flat prior to an optimal *d*_*max*_ as the score is largely determined by *AIC*_*c*_. After *d*_*max*_, the scoring function increases rapidly as the penalty contributions from oscillations, presence of negative values and finish near *d*_*max*_ become more substantial. The SEC-SAXS samples are structurally homogeneous and mono-dispersed implying that each dataset can be described by a single *d*_*max*_ value. For more challenging samples such as the unpurified BSA dataset, the overall score does not show a clear minimum, in fact the likelihood function shows multiple peaks (Figure 6C) implying an ambiguity in assigning *d*_*max*_. The sample is dominated by monomers but also contains dimers and additional oligomers suggesting *d*_*max*_ will not be due to a single structural species. The ambiguity in determining a definitive *d*_*max*_ reflects the quality of the sample and in this regard, the likelihood function illustrates a single, symmetric peak for well defined, mono-dispersed samples versus multiple or broad peaks (Figure 6C and D) for poly-dispersed systems. The search method and use of our score into a likelihood function was further tested on non-globular particles that include a DDM micelle, 25 bp dsDNA with 10 nucleotide single stranded overhang (17), 50 bp dsDNA, unfolded SAM riboswitch in EDTA (26), amphipol stabilized G-protein coupled-receptor (27), and BMV TLS RNA (28) (see Figures S6 and S7).

**Figure 6:**
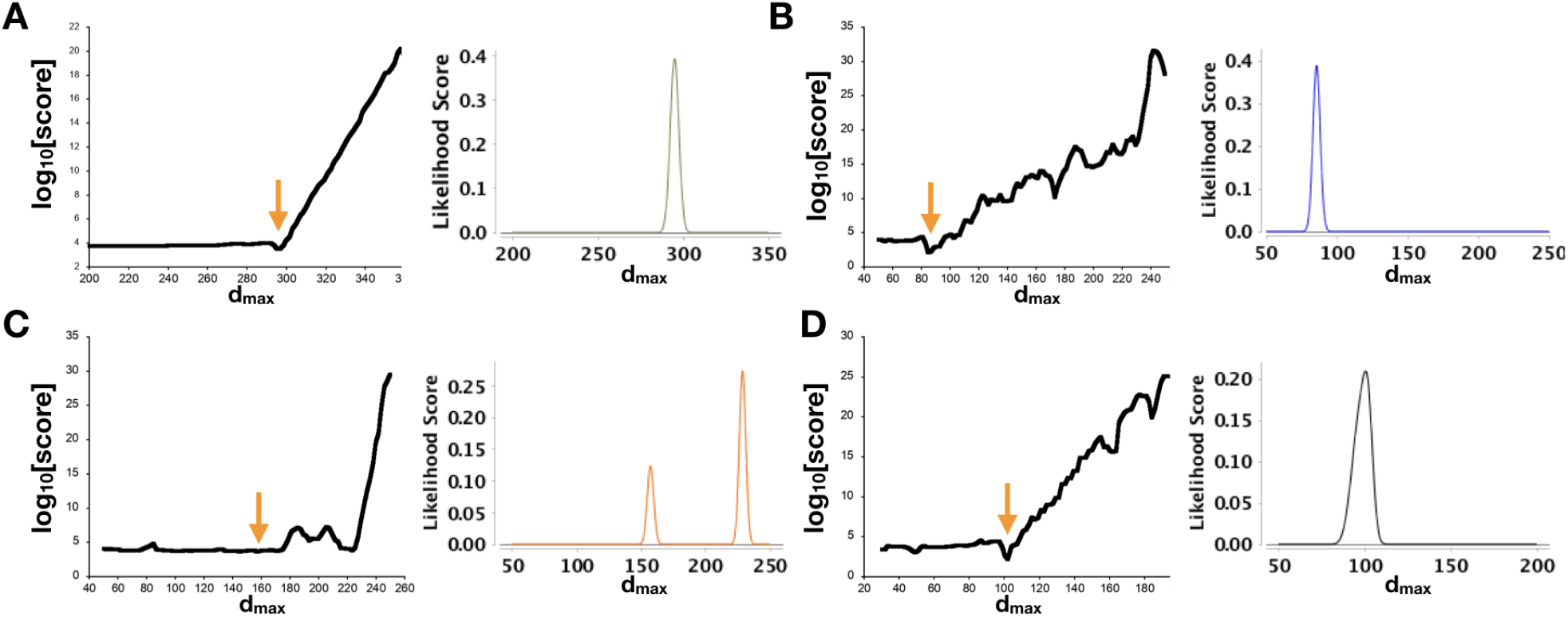
Scoring of *P*(*r*)-distributions in a *α, d*_*max*_ search. A) coil-coiled protein from Figures 2D. B) SEC-SAXS purified BSA from Figure 2A. C) Unpurified BSA from Figure 4D. D) Mixture of BMV TLS RNA (*d*_*max*_ 108 Å) and xylanase (*d*_*max*_ 43 Å). Note the asymmetry in the likelihood peak or presence of multiple peaks for samples that are mixtures. For each panel, the righthand plot illustrates the likelihood score calculated from the set of scores for each search. All searches were performed with the Moore method using smoothness regularization and all x-axis are in units of Å. The *d*_*max*_ search range is given by the x-axis for each panel. BMV TLS RNA was kindly provided by Jeff Kieft, University Colorado School of Medicine, Boulder, CO.

## DISCUSSION

Since the foundational contributions of Glatter and Moore in the 1970s, the modern advances in X-ray focusing optics, monochromators and detector technologies now provide highly over-sampled and focused SAXS measurements that are free of smearing artefacts (10, 29–31). Likewise, advances in sample preparation including the availability of SEC-SAXS at synchrotron facilities (32), has enabled perfect buffer/background matching for producing the ideal, subtracted SAXS signals of biological particles. This convergence of sample preparation and instrumentation now provides for reliable and routine measurements of the SAXS signal that can be exploited to overcome previous limitations.

Using the information theory framework, we see that the so-called Shannon number, *N*_*s*_, corresponds to the boundary between a well- and ill-conditioned problem. For a given [*q, d*]_*max*_, this boundary position determines the maximum number of orthogonal components required to fully describe the inverse SAXS problem. Any additional components makes the problem ill-conditioned thereby requiring some form of regularization (e.g., non-negativity or smoothness) to solve for the unknowns. A non-negativity regularization reduces the size of the search space whereas a smoothness regularization imposes correlations between *P*(*r*)-values to reduce the variation between neighboring values. Alternatively, an orthogonality regularization could be performed to optimally maintain independence between the unknowns. Nevertheless, regularization is a balance between maintaining strict independence of the unknowns in 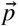 and the orthogonality of the columns of the *S*-matrix. This balance can be achieved best by restricting the complexity of the problem to *N*_*s*_ and utilizing a low *α* in regularization. The goal of the inverse transform is to recover the real-space information that is least biased, free of algorithm or regularization artefacts.

The SPI method solves for a 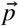with *N*_*s*_ terms, by dividing the *P*(*r*)-distribution into bins of equal width, *b*_*w*_, given by *d*_*max*_/*N*_*s*_. Using the relationship described by Moore, we see that *b*_*w*_ is fundamentally determined by *q*_*max*_ (Eq. 11). Regardless of the size of the particle, the effective resolution of the *P*(*r*)-distribution is essentially *b*_*w*_ meaning that no distance can be resolved to better than *b*_*w*_.

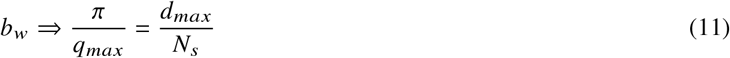

Eq. 11 also implies that a higher resolution measurement, *i*.*e*., increasing *q*_*max*_, will increase *N*_*s*_ thereby reducing the width of the bin used to define the *P*(*r*)-distribution. A smaller *b*_*w*_ increases the level of detail in the *P*(*r*)-distribution producing a higher information content SAXS measurement. This relationship is invariant to how the inverse transform is performed, *e*.*g*., SPI, Legendre, Moore, GNOM, or GIFT. Notably, it is only through increasing *q*_*max*_ that the information content or details of the *P*(*r*)-distribution can be increased.

Inverse integral transform problems using data with Gaussian, white noise will not produce a solution that is uniquely determined with high confidence when using a *χ*^2^-like metric. Various methods have been proposed including Bayesian-based (14) and hybrid scores (9) to assess model-data agreements similar to the approach presented above. Determining a *d*_*max*_ from a bioSAXS experiment is essentially accepting the *d*_*max*_ that gives the most acceptable features in the distribution. Proposed nearly 30 years ago, this perception (9) of the best answer was developed into a score or perceptual criteria that quantified a leading SAXS expert’s experience and knowledge into a useable score for the inexperienced, novice user. Using *AIC*_*c*_, our approach improves upon this type of approach by further considering the complexity of the inverse transform model. Our proposed score (Eq. 10) does have its limitations as it penalizes for negative *P*(*r*)-values. There may be instances in contrast variation or anomalous SAXS measurements that support negative values in the *P*(*r*)-distribution and in these cases, it would be prudent to consider a different method for assessing Ψ. Eq. 10 is modular and readily allows adjustments to Ψ to compensate for atypical SAS experiments. The *AIC*_*c*_ finds the least complex answer that is balanced by perception akin to Occam’s razor.

## CONCLUSION

BioSAXS is the most relevant technique for providing comprehensive measurements of dynamic macromolecular conformations and assemblies in solution that are directly relevant to determining biological structure–function relationships (1, 33, 34). From examination of exemplary modern SAXS data sets, we find that combined advances in synchrotron optics and X-ray data collection provide oversampled datasets suitable for a more complete application of informational theory than has been applied to date. Furthermore, we present quantitative methods to identify models in best agreement with observed scattering data while avoiding over-fitting. The reported methods and findings robustly determine the critical maximum dimension value, provide a quantitative measure of resolution, and objectively test model-data agreement. SAXS routinely extends and complements structural knowledge obtained from crystallography (35), NMR (36–38) and now, cryo-EM (39), where MD-coupled SAXS modeling can discover new biological insights and assist in computational protein design (40, 41). In fact, cryo-EM of macromolecular machines, such as the transcription preinitiation complex (42), is suggesting large motions are fundamental dynamical attributes that are well suited for studies by SAXS. A rigorous direct inverse Fourier transform of the scattering data that assesses both the quality of the recovered-distribution and the information limit (resolution) not only improves the reliability of SAS experiments but also advances insights by minimising over-fitting during the atomistic modelling of SAS. Looking ahead, SAXS has a critical role in understanding functional dynamicity, a concept implying that the regulation and activities of assemblies is achieved through perturbations in their inherent dynamics. SAXS observations can efficiently inform on such dynamicity exemplified by studies investigating impact of mutations, post-translational modifications, activity differences between human versus homologs, and even high-throughput, time-resolved screens for allosteric inhibitors that alter subunit conformations as well as assembly (43–50). We, therefore, expect that these methods and concepts will provide enabling criteria to extend the boundaries for current bioSAS analyses.

## AUTHOR CONTRIBUTIONS

RP Rambo and JA Tainer contributed to the conception and design of the study as well as data analysis and interpretation. RP Rambo developed the theory and mathematical implementations for coding in JAVA. RP Rambo and JA Tainer wrote the manuscript.

## ACKNOWLEDGMENTS

We thank N. Cowieson, K. Inuoue and Nikul Khunti of Diamond Light Source (Didcot, UK), J. Doutch (ISIS, Harwell UK), O. Davies, and M. Tully (ESRF, FR) for helpful discussions and access to datasets. We acknowledgment the SIBYLS beamline (Advanced Light Source, Berkeley CA) and B21 (Diamond Light Source limited, Didcot, UK) for data collection time. J.A.T. acknowledges support by NIH R35 CA220430, NIH P30 GM124169, and Robert A. Welch Chemistry Chair. The SIBYLS synchrotron beamline 12.3.1 and initial efforts by R.P.R. were supported by the Department of Energy, Office of Basic Energy Sciences, Integrated Diffraction Analysis Technologies (IDAT) program.

## SUPPLEMENTARY MATERIAL

An online supplement to this article can be found by visiting BJ Online at http://www.biophysj.org.

## Notes

### Competing Interest Statement

The authors have declared no competing interest.

