## Supporting Information for "Information Theory Optimization of Signals from Small Angle Scattering Measurements"

### 1 Supporting Information

#### 1.1 Presentation of the Moore sine integral transform method

SAXS of dilute, non-interacting homogeneous particles in solution is described by the Debye equation that relates, at a given $q_j$, the distribution of internal distances to an observed $I(q_j)$ through an integral transform.

$$I(q_j) = 4\pi \int P(r)\, \frac{sin(q_j r)}{q_j r}\, dr \tag{1}$$

To put (1) in standard form[10], we rearrange terms $q$ and $r$ as

$$q_j \cdot I(q_j) = 4\pi \int \frac{P(r)}{r}\, sin(q_j r)\, dr \tag{2}$$

and by letting $Q(r) = P(r)/r$ and $U(q) = q \times I(q)$ and substituting yields

$$U(q_j) = 4\pi \int Q(r)\, sin(q_j r)\, dr \tag{3}$$

This clearly demonstrates that the relationship between real, $Q$, and reciprocal space, $U$, is through a sine-integral transform. $Q(r)$ is an odd function since $Q(-r)$ must be equal to $Q(r)$. $Q(r)$ can be represented by any orthogonal series expansion compatible with odd functions such as the Fourier sine series, Chebyshev or Legendre polynomials [11, 8]. Using the Fourier sine series, Moore showed the integral in (3) is a linear function with unknowns, $a_m$.

$$U(q_j) = 2\, d_{max} \sum_{m=1}^{N_s} a_m \cdot m \cdot (-1)^{m+1} \frac{sin(q_j d_{max})}{(m\pi)^2 - (q_j d_{max})^2} \tag{4}$$

In matrix form, the $\vec{p}$ will consist of the Moore coefficients elements, $a_m$, for $m$ in $\{1 \ldots N_s\}$. The $S$-matrix will contain $J$ rows (one row for each data point at $q_j$) and $N_s$ columns. Each element of the $S$-matrix will contain a single term from the series calculated at a specified $q_j$ and $m$ value.

$$\begin{bmatrix} 1 \cdot (-1)^{1+1} \frac{sin(q_1 d_{max})}{(1 \cdot \pi)^2 - (q_1 d_{max})^2} & \cdots & N_s \cdot (-1)^{N_s+1} \frac{sin(q_1 d_{max})}{(N_s \pi)^2 - (q_1 d_{max})^2} \\ \vdots & \vdots & \vdots \\ \vdots & \vdots & \vdots \\ 1 \cdot (-1)^{1+1} \frac{sin(q_J d_{max})}{(1 \cdot \pi)^2 - (q_J d_{max})^2} & \cdots & N_s \cdot (-1)^{N_s+1} \frac{sin(q_J d_{max})}{(N_s \pi)^2 - (q_J d_{max})^2} \end{bmatrix} \cdot \begin{bmatrix} a_1 \\ \vdots \\ \vdots \\ a_{N_s} \end{bmatrix} = \begin{bmatrix} q_1 \cdot I(q_1) \\ \vdots \\ \vdots \\ q_J \cdot I(q_J) \end{bmatrix} \tag{5}$$

Therefore, the unknowns in the $\vec{p}$ can be recovered through an inversion of the $S$-matrix using a suitable pseudo-inverse method. In the presence of constant background, an additional column of ones and element will be added to the $S$-matrix and $\vec{p}$, respectively.

$$\begin{bmatrix} 1 & 1 \cdot (-1)^{1+1} \frac{sin(q_1 d_{max})}{(1 \cdot \pi)^2 - (q_1 d_{max})^2} & \cdots & N_s \cdot (-1)^{N_s+1} \frac{sin(q_1 d_{max})}{(N_s \pi)^2 - (q_1 d_{max})^2} \\ \vdots & \vdots & \vdots & \vdots \\ \vdots & \vdots & \vdots & \vdots \\ 1 & 1 \cdot (-1)^{1+1} \frac{sin(q_J d_{max})}{(1 \cdot \pi)^2 - (q_J d_{max})^2} & \cdots & N_s \cdot (-1)^{N_s+1} \frac{sin(q_J d_{max})}{(N_s \pi)^2 - (q_J d_{max})^2} \end{bmatrix} \cdot \begin{bmatrix} a_0 \\ a_1 \\ \vdots \\ a_{N_s} \end{bmatrix} = \begin{bmatrix} q_1 \cdot I(q_1) \\ \vdots \\ \vdots \\ q_J \cdot I(q_J) \end{bmatrix} \tag{6}$$

The Moore coefficients can be used to calculate the $P(r)$-distribution through Eqn. 7.

$$P(r) = \frac{1}{2\pi^2} \cdot r \cdot \sum_{m=1}^{N_s} a_m \, sin\left(\frac{m \cdot \pi \cdot r}{d_{max}}\right) \tag{7}$$

#### 1.2 Presentation of the Legendre orthogonal series expansion

Similar to the Moore method, the Legendre series expansion approximates [11] the intensity data as $q \times I(q)$ with the exception that we are only expanding the unknown distribution and not $P(r)/r$. Here, the density function, $P(r)$, is strictly defined within the interval [-1,1] and can be approximated using a Legendre polynomial expansion as

$$P(r) = \sum_{m=0}^{\infty} \lambda_m \, L_m\left(\frac{2 \cdot r - d_{max}}{d_{max}}\right) \tag{8}$$

The expansion is infinite but we will restrict the approximation to $N_s$. The evaluation of Legendre polynomial of order $m$, $L_m$, occurs at specified $r$-values that define the $P(r)$-distribution calculated as half-integer increments of the bin-width $(b_w)$

$$b_w = \frac{d_{max}}{N_s} \Rightarrow r_i = (0.5 + i) \cdot b_w \; where \; i \in \{0 \cdots N_s\} \tag{9}$$

$N_s$ is taken as the ceiling of the product of $(q_{max} \cdot d_{max} \cdot \pi^{-1})$. The matrix representation using the Legendre polynomials is given by

$$\begin{bmatrix} \sum \frac{sin(q_1 r)}{r} \cdot L_0\left(\frac{2 \cdot r - d_{max}}{d_{max}}\right) & \cdots & \sum \frac{sin(q_1 r)}{r} \cdot L_{N_s}\left(\frac{2 \cdot r - d_{max}}{d_{max}}\right) \\ \vdots & \vdots & \vdots \\ \vdots & \vdots & \vdots \\ \sum \frac{sin(q_J r)}{r} \cdot L_0\left(\frac{2 \cdot r - d_{max}}{d_{max}}\right) & \cdots & \sum \frac{sin(q_J r)}{r} \cdot L_{N_s}\left(\frac{2 \cdot r - d_{max}}{d_{max}}\right) \end{bmatrix} \cdot \begin{bmatrix} \lambda_0 \\ \lambda_1 \\ \vdots \\ \lambda_{N_s} \end{bmatrix} = \begin{bmatrix} q_1 \cdot I(q_1) \\ \vdots \\ \vdots \\ q_J \cdot I(q_J) \end{bmatrix} \tag{10}$$

which shows the $\vec{p}$ contains the unknown Legendre coefficients, $\lambda_m$, and the $S$-matrix contains product-sum terms where each element is a sum over all $r$-values involving the product of a Legendre polynomial of fixed order and a sine term. The $\vec{p}$ can be recovered using a suitable matrix inversion technique.

#### 1.3 Moore Method with Regularization

The Moore method determines a set of unknown coefficients that describe both the intensity data (Eqn. 4) and the real-space $P(r)$-distribution (Eqn. 7). We can use Eqn. 4 in a regularization scheme where the canonical objective function in least squares is augmented (Eqn. 11) with the L2-norm of the second derivative of the $P(r)$-distribution (Eqn. 7).

$$minimize: \; \|S \cdot \vec{p} - U(\vec{q})\|^2 + \alpha \cdot \left\| \frac{d^2}{dr^2} P(r) \right\|_2 \tag{11}$$

The L2-norm of the second derivative acts as a smoothness constraint and is strictly positive[6, 14]. A perfectly smooth function will have a slope of zero representing a horizontal line.

$$\left\| \frac{d^2}{dr^2} P(r) \right\|_2 = \sum_{i=1}^{r\ values} \left( \frac{d^2}{dr^2} P(r_i) \right)^2 \tag{12}$$

Eqn. 7 is a closed-form, twice differentiable function that allows direct calculation of the second derivative at each $r$-value used in the $P(r)$-distribution (Eqn. 13).

$$\frac{d^2}{dr^2} P(r) = \frac{1}{\pi \cdot d_{max}} \sum_{m=1}^{N_s} a_m \cdot m \cdot cos\left( \frac{m \cdot \pi \cdot r}{d_{max}} \right) - r_i \cdot \frac{1}{\pi \cdot d_{max}^2} \sum_{m=1}^{N_s} a_m \cdot m^2 \cdot sin\left( \frac{m \cdot \pi \cdot r}{d_{max}} \right) \tag{13}$$

The analytical expression for the second derivative of the $P(r)$-distribution given by the Moore equation provides for an expression of the gradient, $\nabla_{a_m}$, of Eqn 11 with respect to the unknown coefficients. The unknowns can be solved using any suitable minimizer or directly through a matrix factorization technique.

$$\nabla_{a_m} \to -2 \cdot \left( S \cdot \vec{p} - \vec{U}(q) \right) \cdot \left[ 2d_{max} \cdot m \cdot (-1)^{m+1} \sum_{j}^{q-values} \frac{sin(q_j d_{max})}{(m\pi)^2 - (q_j d_{max})^2} \right] + \alpha \cdot \frac{d}{da_m} \cdot \frac{d^2}{dr^2} P(r) \tag{14}$$

#### 1.4 Scoring Function

Determining an acceptable $P(r)$-distribution from the SAXS dataset can not be achieved reliably through a simple, residuals-based $\chi^2$-like metric alone. Such a statistic is prone to over-fitting and does not account for the correlations that exist in a SAXS dataset[12]. Often, the acceptability of a real-space transform is achieved by the user where the quality of the $P(r)$-distribution is subjectively evaluated for smoothness, negative values and oscillations. Such a set of expectations was nicely quantified by Svergun[14] into a set of perceptual criteria and implemented in the program GNOM[14]. In our implementation, we recognize that a search is initially performed over some finite $d_{max}$ space. For $d_{max}$ values that are too short, $\chi^2$-like metrics tend to be good enough to recognize poorly fitting models (large residual discrepancies), however, when $d_{max}$ values are too large, $\chi^2$-like metrics become less discriminating as the increased model complexity produces small variations in the residuals. To compensate for this, we utilize the Akaike Information Criteria, AIC[1]. AIC requires an estimate of the model's number of parameters, $k$, and likelihood score used to evaluate the model-data agreement. For low sample sizes, $n$, an additional term is required to give a sample size corrected $\mathrm{AIC_c}$ (Eqn. 13)[3]

$$AIC = 2 \cdot k - 2 \cdot ln(L) + \frac{2k \cdot (k+1)}{n - k + 1} \tag{15}$$

We can approximate the likelihood score, L, (Eqn. 16)using a residual-based metric such as $\chi^2$.

$$L = \prod_{j}^{obs} \frac{1}{\sqrt{2\pi \cdot \sigma_j^2}} \, e^{-\frac{(I(q_j)_{calc} - I(q_j)_{obs})^2}{2\,\sigma_j^2}} \approx e^{-\chi^2} \tag{16}$$

To minimize the effects of the correlations in the SAXS dataset, we choose $\chi^2_{free}$ as our residuals-based metric. $\chi^2_{free}$ is normalized in our program; therefore, we multiply by $N_s$ to rescale. The number of parameters of the model, $k$, must include considerations regarding the choice of $\alpha$, $d_{max}$ and model (*i.e.*, Moore, SPI, or Legendre method), thus we define the total number of parameters, $d_f$, as $N_s + 3$. Finally, the total number of points or sample size used in the $\chi^2_{free}$ calculation is given by T. The $AIC_c$ presented in the paper is given by Eqn. 17.

$$AIC_c = 2 \cdot d_f + N_s \cdot \chi^2_{free} + \frac{2d_f \cdot (d_f + 1)}{T - d_f - 1} \tag{17}$$

To provide additional discriminating power to our score, we incorporate the Durbin-Watson ($D_W$) statistic [4](equation 18) which evaluates the randomness of the residuals calculated over all the data points, $J$, defined by $[q_{min},\ q_{max}]$.

$$D_w = \frac{\sum_{j=2}^{J}(R_j \cdot R_{j-1})}{\sum_{j=1}^{J} R_j^2} \quad where \;\; R_j = q_j \cdot (I(q_j)^{calc} - I(q_j)^{exp}) \tag{18}$$

The $D_W$statistic is an autocorrelation calculated with a lag of 1. Ideally random residuals will have a $D_W$ of 2 and is bounded between 0 and 4. Since we are seeking a minimum for our scoring function, we will use the absolute value of (2 - $D_W$) to inform on the quality of model-data agreement over the entire SAXS dataset defined by $[q_{min},\ q_{max}]$.

Finally, to assess the quality of the $P(r)$-distribution, we quantify smoothness, presence of negative values and oscillations. Smoothness is calculated as the totality of two different features: 1) the absolute sum of the second derivative calculated at the points that define the distribution function and 2) the finish near $d_{max}$. The second derivative sum is normalized to the integrated area of the distribution. To assess the finish at $d_{max}$ , we recognize that $P(r)$-distributions calculated from atomistic coordinates are exceptionally shallow, the finish is free of bumps or bulges. Rather than calculating the slope, which will scale with the size of the particle or data normalization differences, we quantitate the finish based on the angle. The $P(r)$-distribution is centered at $d_{max}$ by subtracting $d_{max}$ from the set of $r$-values. For each of the last 3 points of the centered distribution, an angle is calculated from the point at $P(r_{N-i})$ for $i$ in {1,2,3} to the origin. The angle is calculated in degrees and we seek to minimize the sum of the angles.

$$smoothness \Rightarrow \frac{10}{\left|log_{10} \sum \frac{d^2}{dr^2} P(r)\right|} + 2 \cdot slope\_score \tag{19}$$

We use $log_{10}$the of the $P(r)$-distribution score since the sum of the second derivative can produce exceedingly small values. Consider second-derivative scores of 3.1x10$^{-6}$ versus 7.8x10$^{-7}$, which will translate to 1.8 and 1.6 respectively using the first term in Eqn. 19.

To assess oscillations, we examine the bin heights for the points after the maximum of the $P(r)$-distribution. These points are examined at the Shannon limit, specifically at $r$-values defined by $b_w$. If there are $N$ values in the distribution, indexed from $i = 0 \cdots (N-1)$, then $r_{N-1} = d_{max}$. We define an oscillation by noting that the last point before $d_{max}$ or $r_{N-2}$ should be less than the previous point, $P(r_{N-3})$ and will always be greater than 0 since $P(d_{max}) = 0$. Therefore, an oscillation will be seen as alternating heights at $r$-values indexed at odd integers, $f$ in $\{3,\ 5,\ 7,\ \ldots\}$, where $P(r_{N-f}) < P(r_{N-f+1})$ and $P(r_{N-f}) < P(r_{N-f-1})$. The oscillations must persist through consecutive numbers within $f$.

The presence of negative values is scored by counting the number of $P(r)$ points, $x$, that are less than 1 but penalizing more severely if the last point before $d_{max}$ is negative. We appreciate that some particles, such as detergent micelles, may demonstrate negative values within the body of the distribution and these are penalized least by only considering negative values after the peak of the distribution. This can be succinctly represented by a ternary expression in Eqn. 20.

$$penalty_{neg} = (P(r_{N-2}) < 0)\ ?\ (19.33 \cdot 7^x)\ or\ (1.933 \cdot 7^x) \tag{20}$$

**Algorithm 1** Penalizing oscillations

$f \leftarrow 0$
$N \leftarrow total\ data\ points$
$total\_after\_peak \leftarrow N - index\_of\_peak$
$continue\_on \leftarrow true$
$penalty \leftarrow 0$
**while** $f < total\_after\_peak$ AND $continue\_on$ **do**
  **if** $P(r_{N-f}) < P(r_{N-f+1})$ AND $P(r_{N-f}) < P(r_{N-f-1})$ **then**
    $diff \leftarrow min(P(r_{N-f+1}) - P(r_{N-f}), P(r_{N-f-1}) - P(r_{N-f}))$
    $penalty \leftarrow penalty + 100 * diff/P(r_{N-f})$
    $f \leftarrow f + 2$
  **else**
    $continue\_on \leftarrow false$
  **end if**
**end while**

Our overall P(r)-distribution score that we seek to minimize is given by Eqn21.

$$Pr\_score = \left( \frac{10}{\left| log_{10} \sum \frac{d^2}{dr^2} P(r) \right|} + 2 \cdot slope\_score \right) + penalty_{osc} + penalty_{neg} \tag{21}$$

#### 1.5 Implementation in Java

The SPI, Moore and Legendre methods are implemented in the JAVA program Scatter available at www.bioisis.net and available on GITHUB at https://github.com/rambor/scatter3. The SPI method with the L1-norm is based on a Matlab implementation[7]. Both the Moore and Legendre methods have been implemented with the L2-norm and make use of the Apache Math Commons library (https://commons.apache.org/proper/commons-math/). In all cases, setting α to zero solves the unregularized problem.

In the Scatter project, under the source directory src/version3/InverseTransform, will be the JAVA classes for the methods: SPI (class SVD), SPI with L1-norm (class SineIntegralTransform), Legendre (class LegendreTransform) and Moore (class MooreTransformApache). The Legendre and Moore methods use an instance of the Apache Math Commons solver founded within the nested JAVA class Linear Solver in each respective class. The Linear Solver classes requires explicity calculation of the gradient as the solver performs a step-wise minimization of the target function. The Linear Implementation of Eqn. 21 can be found in the IndirectFT.java class under the method scoreDistribution.

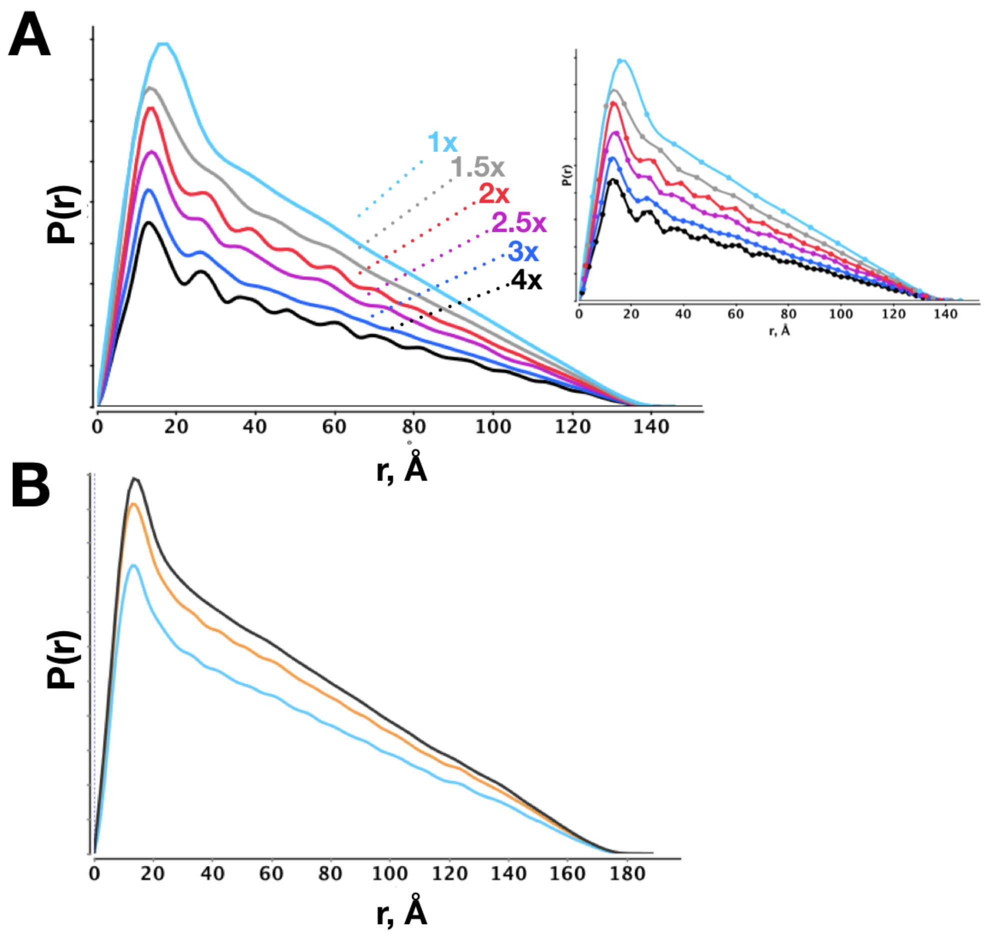


**Figure S1:** $P(r)$-distributions calculated from crystal structures at decreasing bin-widths. A) 137 base-paired double-stranded DNA structure. Bin-widths are calculated at multiples of the B-form helix pitch at k·π/(10.4 Å) for k:{1, 1.5, 2, 2.5, 3, 4} representing $q_{max}$ values of 0.302, 0.453, 0.604, 0.755, 0.906, and 1.208 $Å^{-1}$. Inset shows the actual points (bin heights) determined at each resolution. B) Coiled-coil protein (PDB 5XG2) calculated at $q_{max}$ values of 0.4 (black), 0.6 (orange) and 0.8 (cyan) $Å^{-1}$ representing relative increments of 1x, 1.5x and 2x. Increasing $q_{max}$ shows oscillatory features that are directly derived from the structure. Curves are offset for illustrative purposes. DNA model was kindly provided by Nathan Cowieson, Diamond Light Source, Didcot,UK.

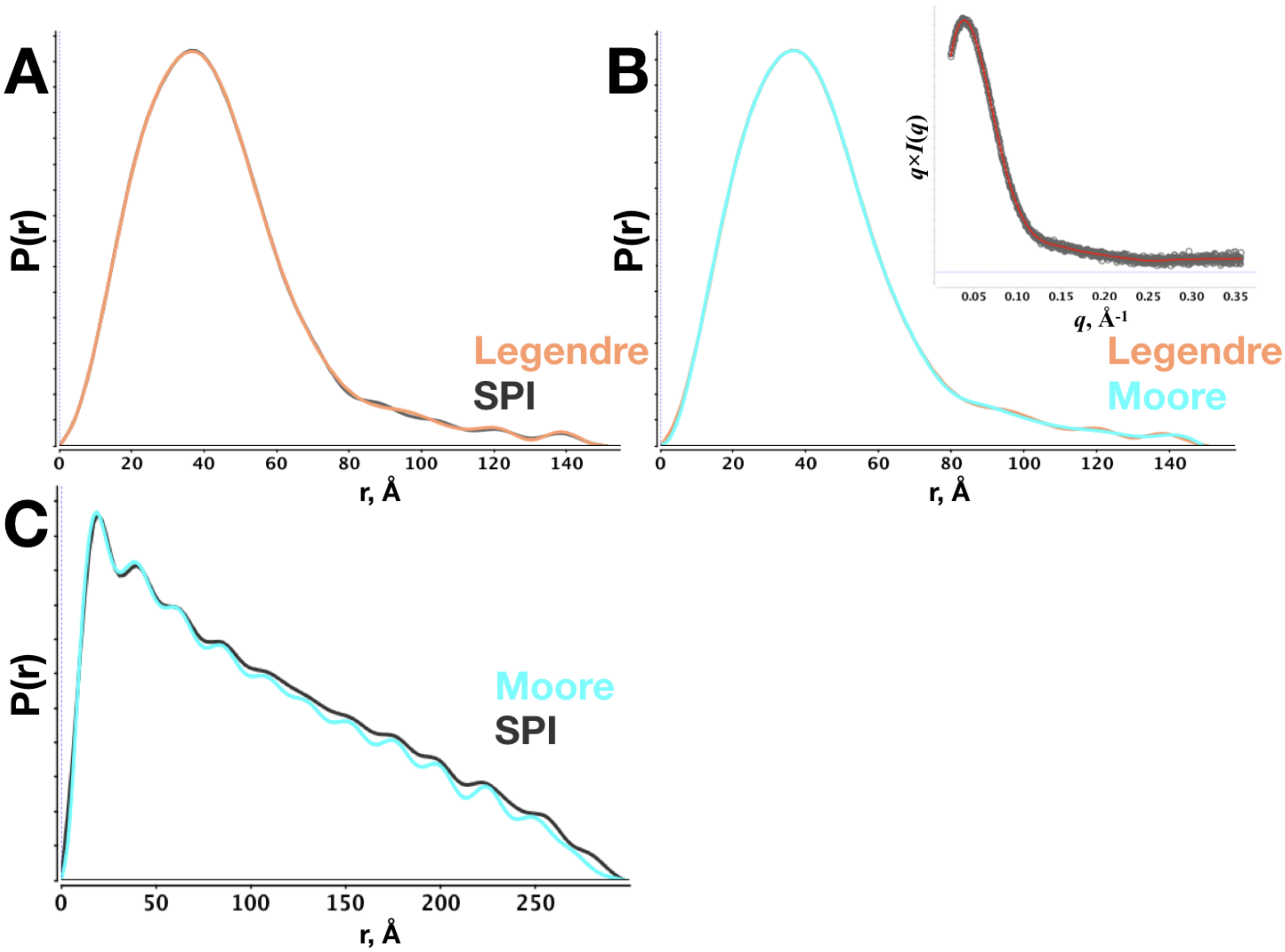


**Figure S2:** Comparison of Legendre, Moore and SPI inverse transform methods without regularization ($\alpha = 0$). A) Overlay of the Legendre polynomial expansion (orange) on the SPI method (gray). B) Overlay of Moore method (cyan) on the Legendre polynomial expansion (orange). Inset shows the SAXS data (black circles) transformed as $q \times I(q)$ with the fit from the Moore method (red line). Data in A and B are un-purified BSA in PBS buffer at 3 mg per mL. C). Overlay of the Moore method (cyan) on the SPI method (gray) using SAXS data of a coiled-coil protein from Figure 2D.

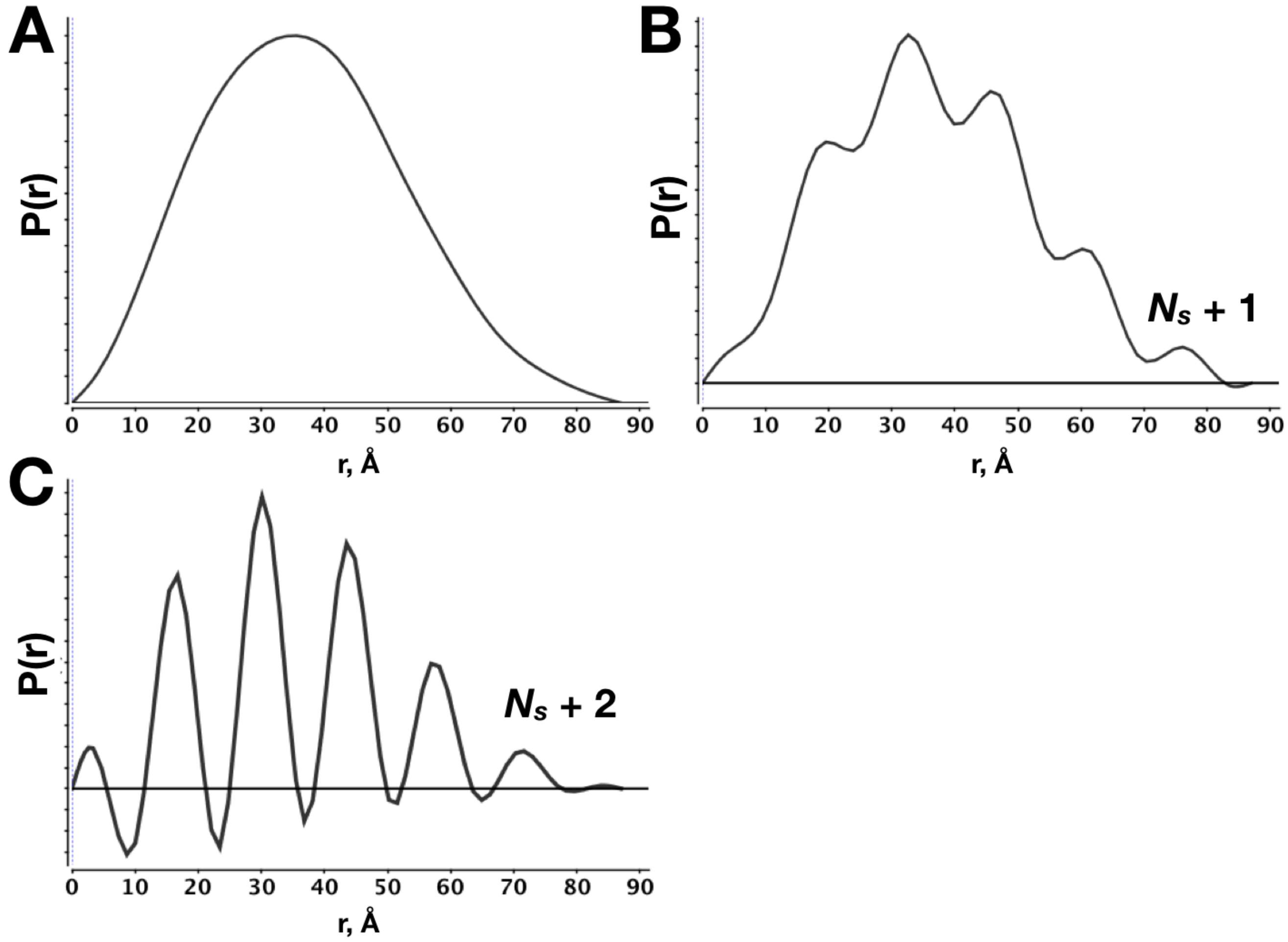


**Figure S3:** Ill-conditioning of the inverse transform as the number of unknowns exceed $N_s$ using the SPI method demonstrated with SEC-SAXS dataset of BSA from Figure 1. A) Transform calculated with $N_s$. B, C Transforms calculated with unknowns exceeding $N_s$.

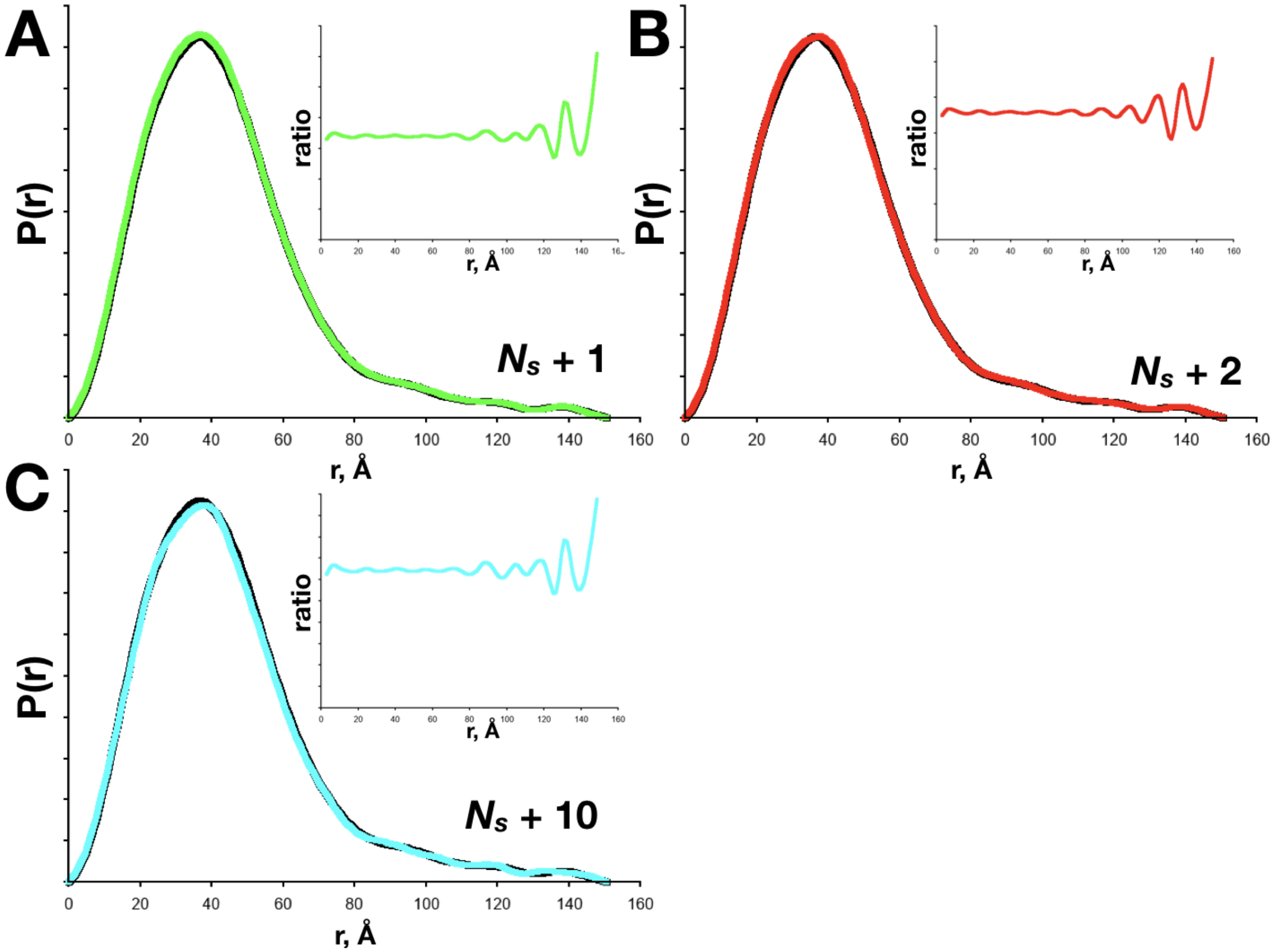


**Figure S4:** Ill-conditioning of the inverse transform as the number of unknowns exceed $N_s$ using an orthogonal series expansion (Legendre polynomials). SEC-SAXS dataset of BSA from Figure 1. Overlay of $P(r)$-distributions calculated using Shannon-limited terms at $N_s$ (black), $N_s$+1 (green), $N_s$+2 (red) and $N_s$+10 (cyan) at a fixed $d_{max}$ of 150.5 without constant background. For each panel, inset illustrates the ratio of each ill-conditioned $P(r)$-distribution to the Shannon-limited $P(r)$-distribution. Undulations present in each ratio show that the distributions are fundamentally different.

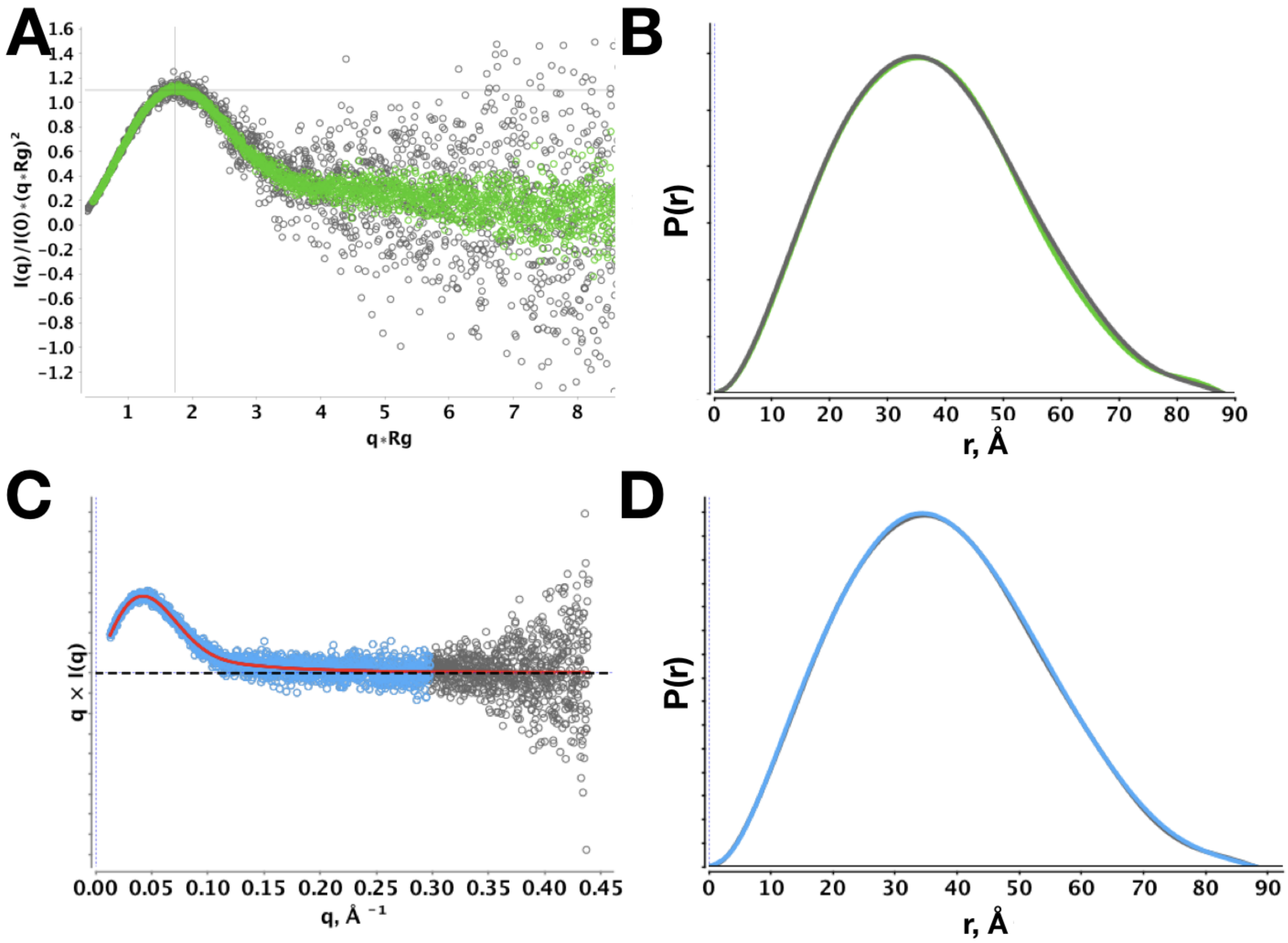


**Figure S5:** Monomeric BSA from SEC-SAXS of dilute BSA in PBS buffer. A) Dimensionless Kratky plot of average of frames across a single elution peak (dark gray circles) corresponding to ≈ 0.26 mg per ml versus average of 18 independent (green), identical SEC-SAXS runs of monomeric BSA. B) Moore-method with L2-norm smoothness regularization for both datasets in A. C) Average of frames across a single elution peak as in A but also truncated (blue circles) to ≈ 0.3 Å$^{-1}$. D) Inverse transform of data in C via Moore-method with smoothness regularization. For both B and D inverse transform was achieved with $\alpha = 331$, $d_{max}$ 90.5 Å.

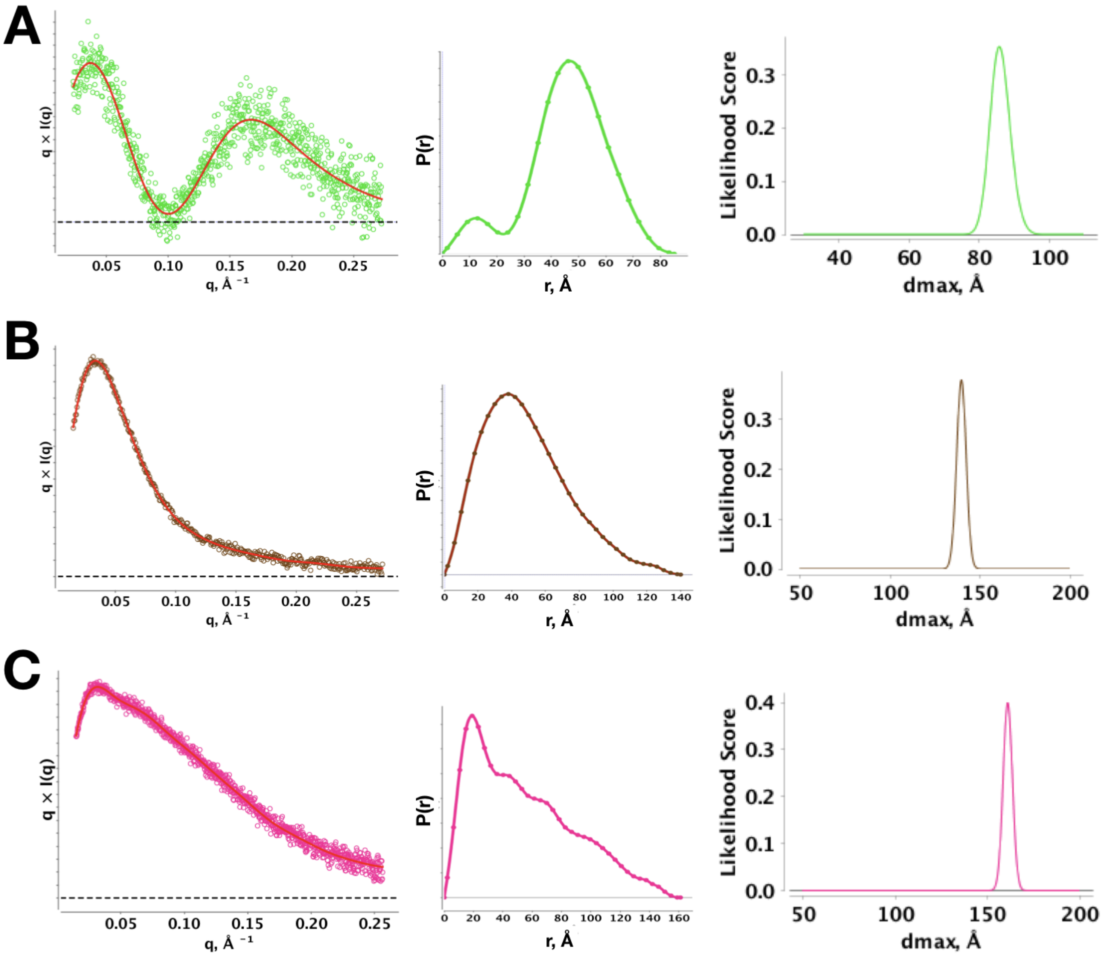


**Figure S6:** Model selection using the Akaike information criteria based score. For each dataset, $d_{max}$ searches were performed within a fixed range as specified by the x-axis of the plots on the right-hand side. The optimal $P(r)$-distribution is plotted along with the score-weighted averaged $P(r)$-distribution. The score-weighted averaged is calculated from the set of distributions contained within the peak(s) (red, central plots). In some cases, the weighted averaged distributions are perfectly overlayed on the single best distribution, colored as in the circles on left-hand plots. A) SEC-SAXS DDM micelle in PBS buffer. B) 25 base-paired, double-stranded DNA with 10 nucleotide single-stranded overhang from [9]. C) SEC-SAXS purified 50 base-paired double-stranded DNA in PBS.

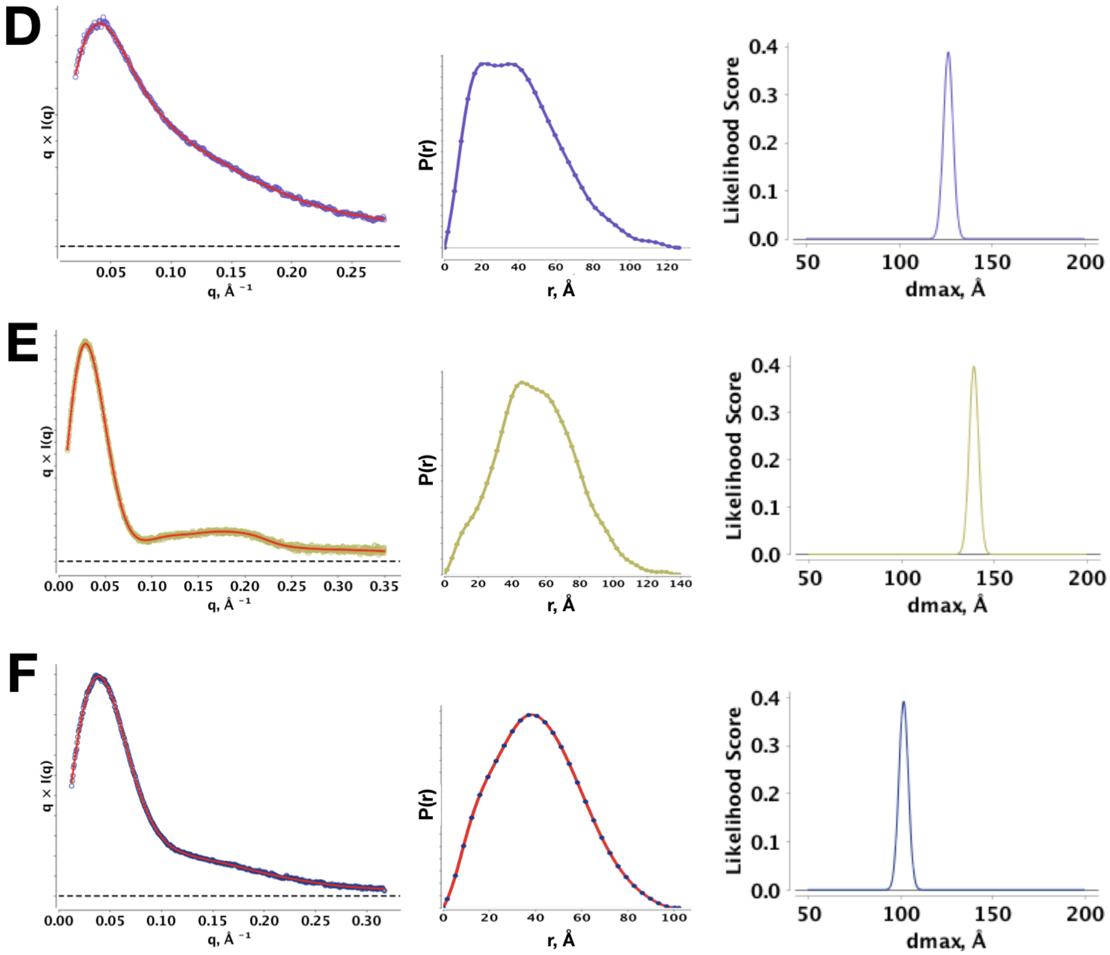


**Figure S7:** Continue of SI Fig. S6 D) unfolded SAM-riboswitch RNA in presence of EDTA from [13]. E) SEC-SAXS of amphipol G-protein coupled receptor from [2]. F) SEC-purified BMV TLS RNA from [5].
